# Connexin 43-mediated Neurovascular Interactions Regulate Neurogenesis in the Adult Brain Subventricular Zone

**DOI:** 10.1101/2022.02.28.482353

**Authors:** Nafiisha Genet, Gael Genet, Jennifer S. Fang, Nicholas W. Chavkin, Hema H. Vasavada, Joshua S. Goldberg, Bipul R. Acharya, Neha Bhatt, Kasey Baker, Stephanie McDonnell, Mahalia R. Huba, Gerry Ma, Anne Eichmann, Jean Leon Thomas, Charles ffrench-Constant, Karen K. Hirschi

## Abstract

The subventricular zone (SVZ) is the largest neural stem cell (NSC) niche in the adult brain; herein, the blood-brain barrier is leaky, allowing direct interactions between NSCs and endothelial cells (ECs). Mechanisms by which direct NSC-EC interactions in the SVZ control NSC behavior are unclear. We found Cx43 to be highly expressed by both NSCs and ECs in the SVZ, and its deletion in either cell type leads to increased NSC proliferation and neuroblast generation, suggesting that Cx43-mediated NSC-EC interactions maintain NSC quiescence. This is further supported by in vitro studies showing co-culture with ECs decreases NSC proliferation and increases their expression of genes associated with quiescence in a Cx43-dependent manner. Cx43 mediates these effects in a channel-independent manner involving its cytoplasmic tail and ERK activation. Such insights further advance our understanding of NSC regulation in vivo and may inform NSC maintenance ex vivo for stem cell therapies for neurodegenerative disorders.

## Introduction

In the adult mouse and human brains, neural stem cells (NSCs) reside in two germinal niches, the dentate gyrus of the hippocampus, or subgranular zone (SGZ), and the subventricular zone (SVZ) of the lateral ventricles, which is the largest NSC niche. In mice, both niches allow the replenishment of new neurons throughout life; the SGZ is involved in hippocampal neurogenesis while the SVZ enables olfactory bulb (OB) neurogenesis. The SVZ is a polarized niche, where neurogenesis is initiated when glial fibrillary acidic protein (GFAP^+^) quiescent NSCs are activated and become epidermal growth factor receptor (EGFR^+^) NSCs that give rise to mammalian achaete-shute homolog 1 (Mash-1^+^) transit-amplifying progenitor cells (TACs)^1–4^. On the ventricular side of the murine SVZ, quiescent NSCs protrude apical processes with short primary cilia that contact the ependymal cell layer lining the ventricles and cerebrospinal fluid within the ventricles^1, 5, 6^. On the parenchymal side, quiescent NSCs project long basal end-feet that make direct contact with blood vessels in the niche^7^.

In the SVZ, TACs assemble into small clusters and differentiate into chains of doublecortin (DCX^+^) neuroblasts^8^; TACs are fast-dividing while neuroblasts proliferate ten times slower^2, 6, 9, 10^. The chains of neuroblasts, along with some clusters of TACs, migrate tangentially from the SVZ through a restricted pathway called the rostral migratory stream (RMS) toward their destination in the OB^2, 11^. Once in the OB, neuroblasts exit the RMS and migrate radially into the granular and periglomerular layers to differentiate into interneurons^9, 12–14^. In the adult SVZ, NSCs predominantly undergo symmetric, differentiative and consuming divisions to generate TACs, which gradually depletes the pool of quiescent NSCs over time^15, 16^. Therefore, it is important to understand the signaling cues between NSCs and the other SVZ niche cells, especially vascular endothelial cells (ECs), to tightly regulate NSC activation, differentiation, migration, and neurogenesis.

It is well-established that SVZ microvascular ECs are fundamental to the niche and support NSC self-renewal, maintenance, proliferation, differentiation, and migration of neural progenitor cells^17, 18^. Several studies have highlighted the role of paracrine signaling between vascular ECs and NSCs in the SVZ via EC-secreted factors such as, vascular endothelial growth factor (VEGF)-A and -C^19, 20^, betacelluline^21^, pigment epithelium-derived factor (PEDF)^22^ and placental growth factor (PlGF) type 2^23^ that regulate NSC behavior in the SVZ. Moreover, in the SVZ microvasculature, and unlike other areas of the brain, the ECs that form the blood vessels lack pericyte and astrocyte coverage, enabling direct contact between NSCs and ECs in this niche^18, 24, 25^. This allows for physical interactions between NSCs and ECs to enable juxtacrine signaling via membrane-bound proteins and their associated ligands to enable Notch and Ephrin signaling^25, 26^. Intercellular junctions, such as gap junctions, also form between NSCs and ECs^27^. Gap junctions enable intercellular signaling via the transfer of ions, metabolites, soluble factors and molecules of ≤1KDa. In the developing mouse brain, gap junction proteins connexin 43 (Cx43) and Cx26 are differentially expressed and play important roles in the regulation of neurogenesis^28, 29^. Gap junctions comprised of Cx43 are known to form between NSCs and ECs in the mouse SVZ and RMS at various postnatal stages^27^. However, the role of EC- or NSC- expressed Cx43 in the regulation of adult SVZ neurogenesis is not known and the focus of this study.

Herein, using an in-vitro transwell co-culture system, we showed that Cx43-mediated interactions between NSCs and ECs decrease NSC proliferation and increase their survival. Furthermore, in adult mice in which *Gja1*, which encodes for Cx43 protein, has been conditionally deleted in either NSCs or ECs for 1 week, there is an initial increase in NSC activation and the generation of DCX^+^ neuroblasts. However, after 4 weeks deletion, there is a significant reduction of quiescent NSCs in the SVZ and increased neurogenesis in the OB. Finally, we show that deletion of *Gja1* in ECs impairs the repopulation of the SVZ niche upon infusion of the antimitotic drug cytosine-β-arabinofuranoside (Ara-C). Collectively, these studies suggest that Cx43-mediated interactions between NSCs and ECs maintain NSC quiescence to regulate neurogenesis in the adult SVZ. Such insights can be applied to the development of ex vivo biomimetic engineered SVZ niches to further study NSC regulation and to potentially use them to treat neurovascular disorders such as stroke.

## Results

### NSC-EC co-culture decreases NSC proliferation in a Cx43-dependent manner

To investigate the effects of ECs on NSC behavior, we used a 2D transwell system where ECs were co-cultured in contact with NSCs allowing for direct cell-to-cell interactions (**Figure 1a and 1b**). In this co-culture model, NSCs were seeded in the absence of EGF and FGF (the growth factors that support NSC survival and growth) to specifically assess the role of ECs in the regulation of NSC behavior. We observed that ECs promote NSC survival, while NSC proliferation measured via EdU incorporation is significantly decreased (**Figure 1c and 1d**). Bulk mRNA sequencing (RNAseq) of NSCs in co-culture with ECs revealed that genes associated with NSC quiescence, such as *Gfap*, *Sox9* and *Prom1,* are upregulated, while genes associated with NSC activation, such as *Egfr and Ccne1,* are downregulated in comparison with NSCs cultured alone (**Figure 1e**). Gene Ontology (GO) analysis revealed that genes associated with neurogenesis are downregulated in NSCs when co-cultured with ECs (**Figure 1f**). Additionally, quantitative (q) PCR analysis performed on NSCs co-cultured with ECs showed increased mRNA levels of *Gfap, Nestin* and *Glast,* genes associated with NSC quiescence while expression of *Egfr, Mash1,* and *Cyclin E,* genes associated with NSC activation, are decreased (**Figure 1g**).

**Figure 1:**
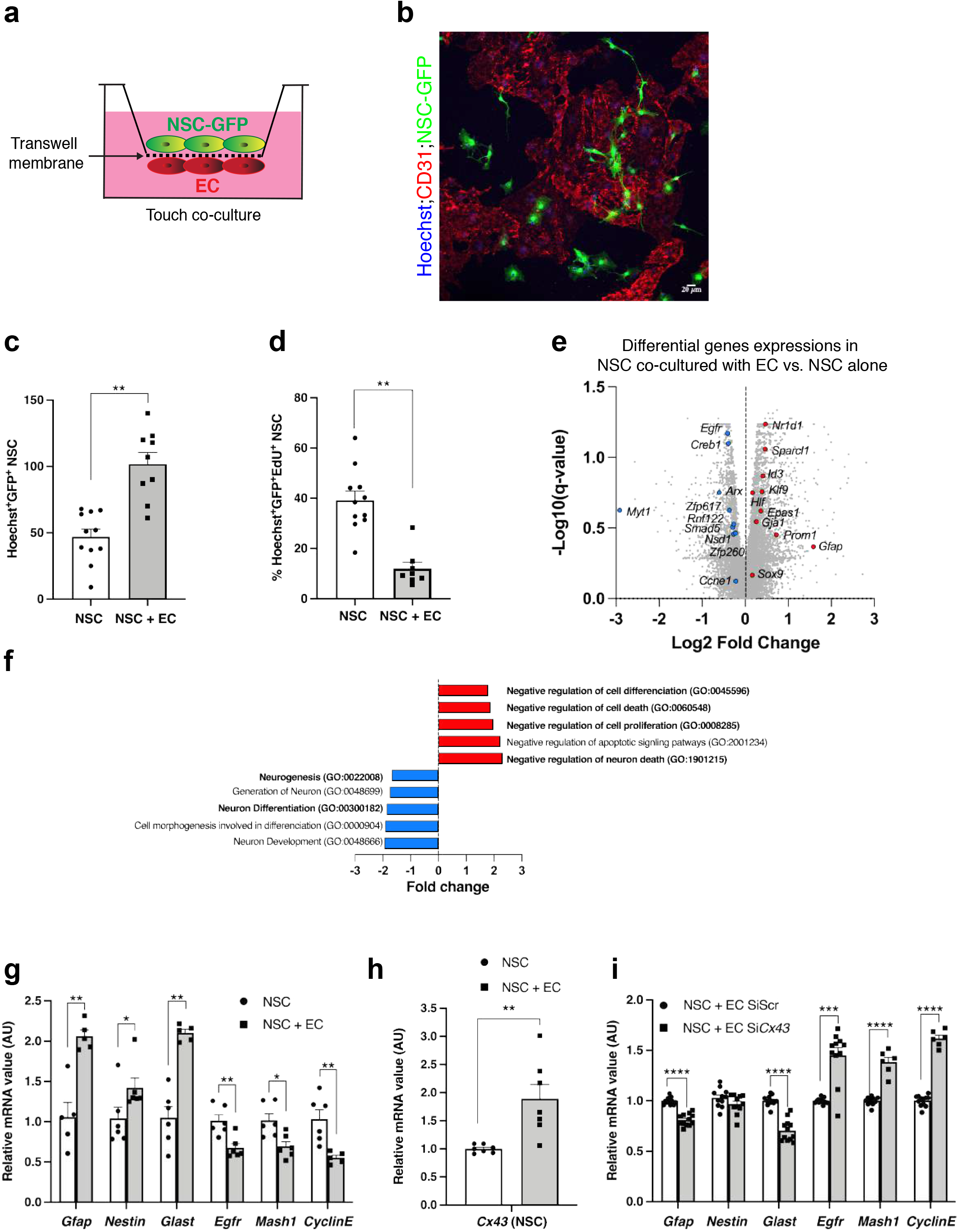
NSC-EC co-culture decreases NSC proliferation in a Cx43-dependent manner. (**a**) Schematic representation for transwell touch co-culture of vascular ECs (bottom side) with GFP^+^NSCs (top side). (**b**) Confocal image showing CD31^+^ EC (red) and GFP^+^ NSCs (green) co-cultured on the transwell membrane. (**c**) Quantification of total GFP^+^NSCs number and (**d**) percentage of GFP^+^NSCs with EdU uptake when cultured alone or with ECs respectively (n=3-4 biological replicates, 4 viewing fields/n). (**e**) Volcano plot of differential genes expressions in NSCs co-cultured with ECs compared to NSCs cultured alone. Red and blue dots indicated selected upregulated or downregulated genes respectively (n=3 biological replicates). (**f**) Gene ontology term analysis of selected gene family modified in NSCs co-cultured with ECs compared to NSCs cultured alone. (**g**) qPCR analysis of quiescent and activated genes in NSCs alone or NSCs co-cultured with ECs (n=3 biological replicates). (**h**) qPCR analysis of *Cx43* gene expression in NSCs alone or NSCs co-cultured with ECs (n=3 biological replicates). (**i**) qPCR analysis of quiescent and activated genes in NSCs co-cultured with ECs, where ECs are treated with control siRNA (siScr) or *Cx43* siRNA (Si*Cx43*) (n=3 different experiments). Data are mean ± SEM. \**p* ≤ 0.05, \*\**p* ≤ 0.01, *** *p* ≤ 0.001.

The RNAseq analysis also revealed the upregulation of potential regulators of NSC-EC interactions (**Figure 1e**), including *Gja1* (also referred to as *Cx43*), which encodes the gap junction protein Cx43. This was of interest to us, as we previously found Cx43 mediates interactions between ECs and vascular mural cells^30^. Thus, we used qPCR analysis to confirm that *Cx43* expression is increased in NSCs when co-cultured with ECs (**Figure 1h**). To determine whether Cx43 plays a role in mediating NSC-EC interactions that lead to changes in NSC gene expression, we used siRNA to suppress *Cx43* expression in ECs and then co-cultured them with NSCs. This resulted in significantly decreased *Gfap* and *Glast* mRNA levels, and significantly upregulated *Egfr, Mash1* and *Cyclin E* mRNA levels (**Figure 1i**). Conversely, when we silenced *Cx43* in NSCs and then co-cultured them with ECs, we did not observe changes in NSC gene expression (**Extended Figure 1a-e**). Collectively, these results suggest that ECs may enhance NSC survival and quiescence via Cx43.

### ECs and NSCs in the adult brain SVZ highly express Cx43

In our RNAseq studies, we found that *Gja1/Cx43* was the only gap junction gene significantly upregulated in NSCs when co-cultured with ECs (**Figure 1e**); however, both cell types have been shown to express other Cx proteins^31, 32^. Thus, we measured the expression of different Cx proteins in ECs and NSCs in vivo in the adult mouse brain SVZ. Using immunohistochemistry and antibodies against different Cx proteins and CD31, which is expressed by all ECs, we found a low percentage of ECs expressing Cx26 and Cx31, while a high percentage of ECs express Cx43 (**Figure 2a**). To evaluate the expression of Cx proteins in NSCs in the murine SVZ, we first labeled the NSCs using a BrdU-labeling protocol in which label-retaining cells (LRC) represent quiescent NSCs^33^. We found that a percentage of SVZ LRC-NSCs express multiple Cx proteins, and a higher percentage of LRC-NSCs express Cx43 (**Figure 2b**), similar to ECs. To determine whether Cx43 is present between NSCs and ECs in the SVZ in vivo, we performed immunohistochemistry and high-resolution confocal imaging of SVZ coronal sections. We found punctate-like Cx43 expression between CD31^+^ ECs and GFAP^+^SOX2^+^ NSCs, as well as between SOX2^+^ cells of the ependymal layer. (**Figure 2c**).

**Figure 2:**
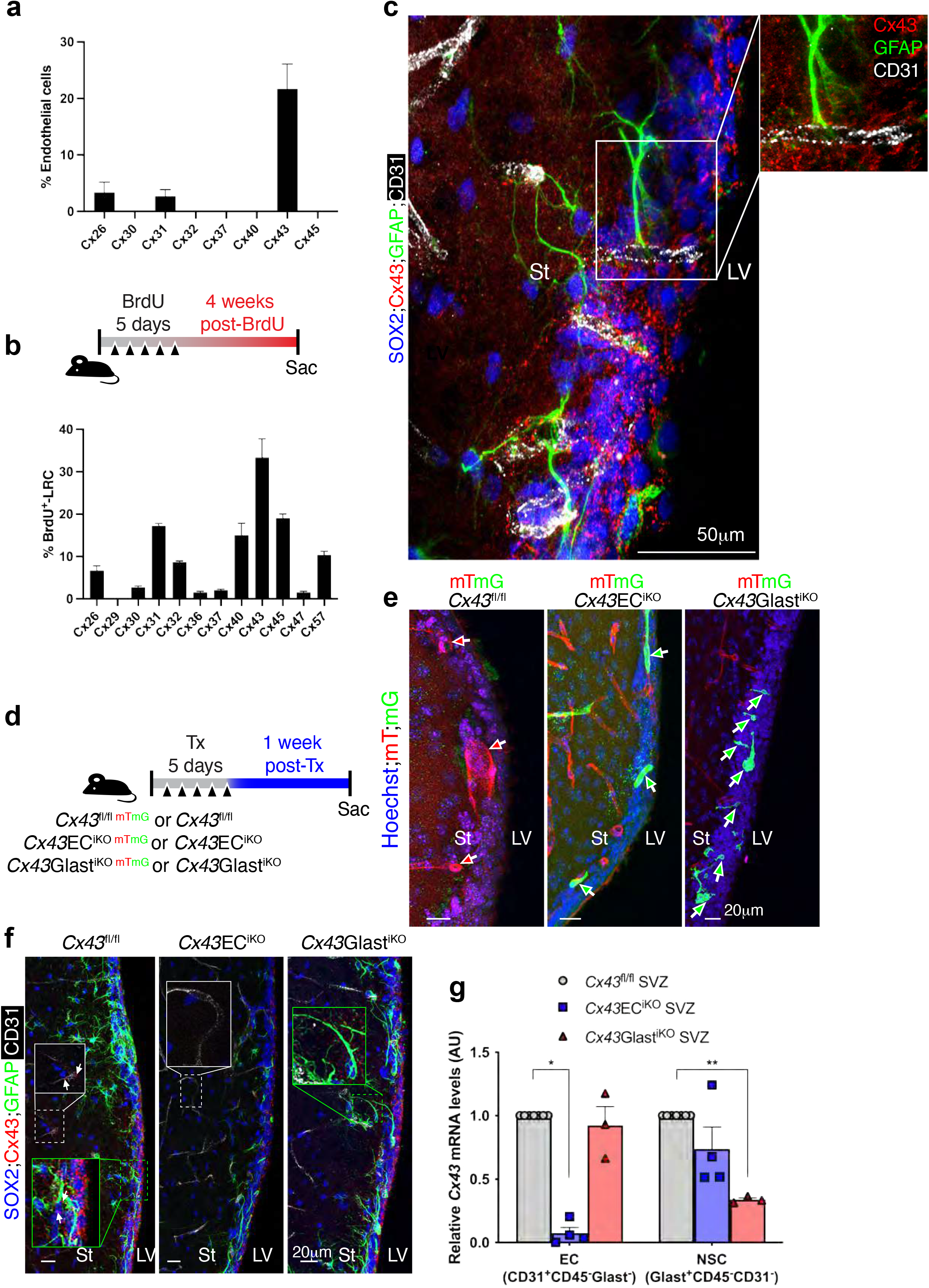
ECs and NSCs in the adult brain SVZ highly express Cx43. (**a**) and (**b**) Quantifications of the percentage of CD31^-^EC and BrdU^+^-LRC^+^ quiescent NSCs colocalizing with different Cx proteins respectively. (**c**) High resolution confocal image of a coronal section from a control mouse brain SVZ showing the end-feet of GFAP^+^SOX2^+^ NSCs (green and blue) contacting CD31^+^ endothelial cells via Cx43 punctate (inset). Scale bar is 50 μm. (**d**) Timeline used to evaluate *Cx43* deletion and recombination in the SVZ. (**e**) Representative confocal images of SVZ from *Cx43*EC^iKO^;ROSA^mT/mG^ and *Cx43*Glast^iKO^;ROSA^mT/mG^ mice. Note the presence of green recombinant ECs or NSCs (green arrows) and only red tomato cells in the *Cx43*^fl/fl^;ROSA^mT/mG^ SVZ (red arrows). (**f**) Representative confocal images of Cx43 immunostaining in the SVZ of *Cx43*^fl/fl^, *Cx43*EC^iKO^ and *Cx43*Glast^iKO^ mice. Note that Cx43 punctate staining (white arrows) in ECs (white inset) and NSC (green inset) are specifically deleted in *Cx43*EC^iKO^ and Cx43Glast^iKO^ SVZ respectively. (**g**) *Cx43* mRNA expression in ECs (CD31^+^CD45^-^Glast^-^ population) and NSCs (Glast^+^CD45^-^CD31^-^ population) of Cx43^fl/fl^, *Cx43*EC^iKO^ and *Cx43*Glast^iKO^ SVZ (n=3-4 different experiments, 5 SVZs per pooled experiment). Scale bar is 20 μm. St: Striatum; LV: Lateral ventricle. Data are mean ± SEM. \**p* ≤ 0.05, \*\**p* ≤ 0.01.

To determine whether NSC- and/or EC-expressed Cx43 plays a role in the regulation of NSCs in the adult brain SVZ, we used *Gja1^flox/flox^* mice (hereafter referred to as *Cx43^fl/fl^* mice) and an inducible loss-of-function genetic approach. To selectively delete *Cx43* in ECs, we crossed *Cx43^fl/fl^* with *Cdh5*Cre^iERT2^ to generate *Cx43*EC^iKO^ mice, and to selectively delete *Cx43* in NSCs, we crossed *Cx43^fl/fl^* mice with *Glast*Cre^iERT2^ to generate *Cx43*Glast^iKO^ mice. To validate Cre-mediated recombination in ECs and NSCs in these models, we crossed both *Cx43*EC^iKO^ and *Cx43*Glast^iKO^ mice with *ROSA^mT/mG^* mice in which, upon tamoxifen (Tx) injection, cell-membrane localized tdTomato converts to GFP in Cre recombinase-expressing cells. We performed recombination and genetic deletion in 6-week-old (adult) mice with Tx injection, as shown in the timeline in **Figure 2d**. As expected, we observed GFP^+^ ECs in the SVZ of *Cx43*EC^iKO^*;ROSA^mT/mG^* mice and GFP^+^ NSCs in the SVZ of *Cx43*Glast^iKO^*;ROSA^mT/mG^* (**Figure 2e**).

To further confirm the loss of Cx43 expression in our mouse models, we also performed immunostaining with antibodies against CD31 to label ECs and antibodies against SOX2 and GFAP to co-label NSCs. In the SVZ of *Cx43*EC^iKO^ mice, we found loss of Cx43 expression in the CD31^+^ ECs and maintenance of Cx43 expression in ependymal cells and NSCs. In the SVZ of *Cx43*Glast^iKO^ mice, we found Cx43 expression was lost in GFAP^+^SOX2^+^ NSCs while ECs and ependymal cells retained expression (**Figure 2f**). *Cx43* deletion efficiency in ECs of *Cx43*EC^iKO^ and NSCs of *Cx43*Glast^iKO^ was confirmed via qPCR analysis of primary SVZ cells dissociated and FACS-isolated into the EC (CD31^+^CD45^-^Glast^-^) and NSC (Glast^+^CD45^-^CD31^-^) fractions (**Extended Figure 2**). Our studies showed that in *Cx43*EC^iKO^ mice, ECs lost ∼90% of *Cx43* expression, which was maintained in NSCs. Conversely, in *Cx43*Glast^iKO^ mice, *Cx43* expression was suppressed by ∼75% in NSCs and maintained in ECs (**Figure 2g**). These results demonstrate effective deletion of Cx43 in SVZ ECs and NSCs in Tx-induced *Cx43*EC^iKO^ and *Cx43*Glast^iKO^ mice, respectively.

### Cx43 deletion in ECs or NSCs depletes quiescent NSCs in the adult SVZ

We next analyzed the consequences of short- (1-week post-Tx) and long-term (4 weeks post-Tx) deletion of *Cx43* in *Cx43*EC^iKO^ and *Cx43*Glast^iKO^ mice, compared to control *Cx43*^fl/fl^, on the number of quiescent and activated NSCs and neuroblasts in the adult SVZ (**Figure 3a and 3b**). The identification of quiescent NSCs is still a challenge due to lack of specific markers. In our study, we used combined expression of the astrocytic marker GFAP and the neural stem and progenitor cell marker SOX2 to identify NSCs ^25, 34, 35^. Since SOX2 and GFAP are expressed by both quiescent and activated NSCs^18^, we injected *Cx43*EC^iKO^ and *Cx43*Glast^iKO^ mice, at 1 week and 4 weeks post-Tx, with EdU 24 hr prior to sacrifice to measure quiescent NSCs (EdU^-^) and activated NSCs (EdU).

**Figure 3:**
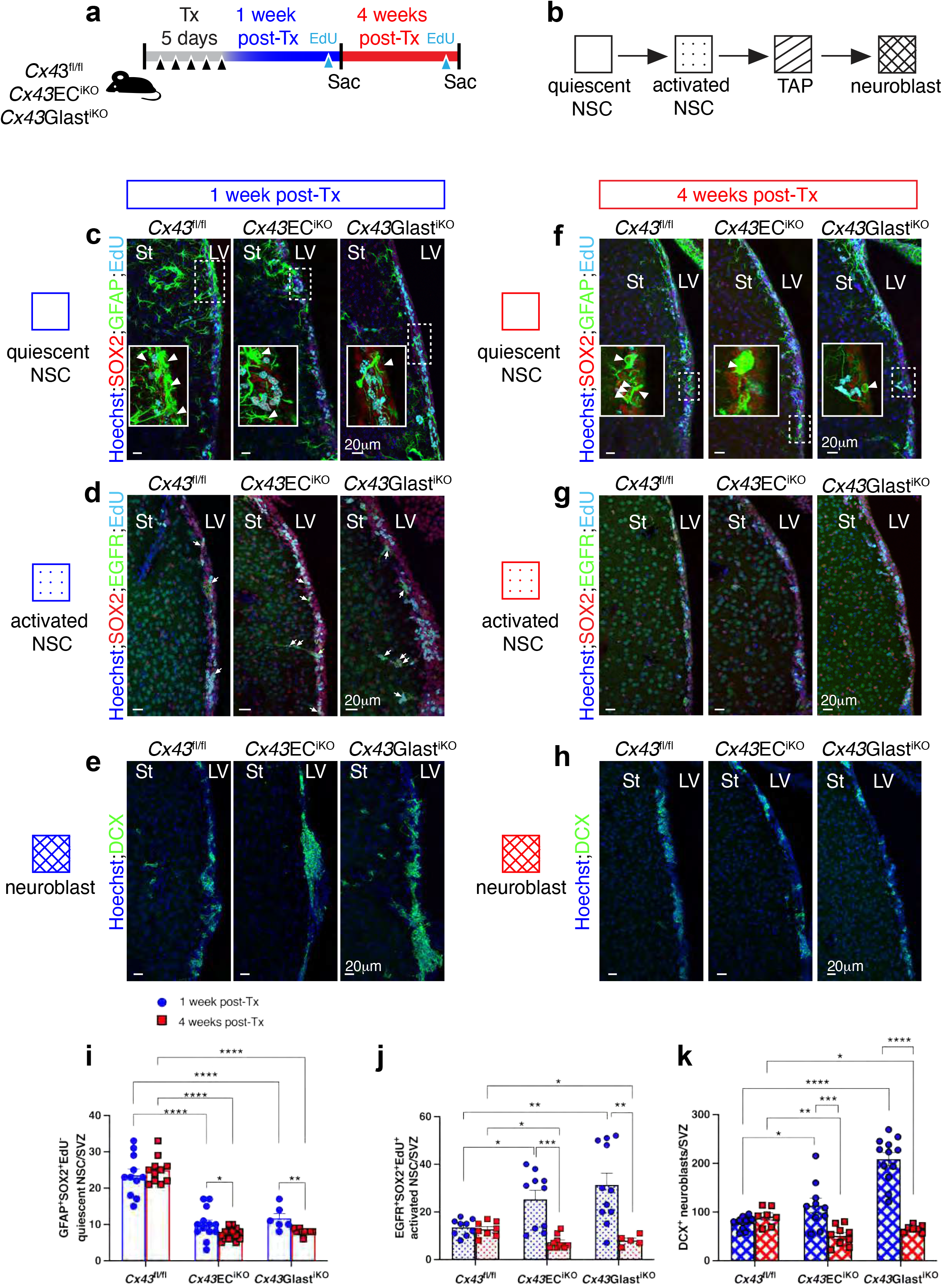
Cx43 deletion in ECs or NSCs depletes quiescent NSCs in the adult SVZ. (**a**) Timeline used to evaluate the effects of short-term deletion (1-week post-Tx) vs long-term deletion (4-weeks post-Tx) of *Cx43* on the number of quiescent NSCs, activated NSCs and neuroblasts in the SVZ. (**b**) Schematic model of quiescent NSCs activation and differentiation in the SVZ. (**c**) to (**h**) Confocal images of SVZ coronal sections from *Cx43*^fl/fl^, *Cx43*EC^iKO^ and *Cx43*Glast^iKO^ mice showing in (**c**) and (**f**) quiescent GFAP^+^SOX2^+^EdU^-^ NSC (arrowheads), (**d**) and (**g**) EGFR^+^SOX2^+^EdU^+^ activated NSCs (arrows in **d**); (**e**) and (**h**) DCX^+^ neuroblasts. (**i**) to (**k**) Quantifications of images shown in (c) to (h) (n=3-5 different animals per group). Scale bar is 20 μm. St: Striatum; LV: Lateral ventricle. Data are mean ± SEM. \**p* ≤ 0.05, \*\**p* ≤ 0.01, \*\*\*\**p* ≤ 0.001.

At 1-week post-*Cx43* deletion, we observed a decrease in the number of GFAP^+^SOX2^+^EdU^-^ quiescent NSCs in both *Cx43*EC^iKO^ and *Cx43*Glast^iKO^ SVZ (**Figure 3c and 3i**) and further reduction at 4 weeks post-Tx (**Figure 3f and 3i**). Additionally, the absence of GFAP^+^ cells co-expressing S100β in the striatum of the mutant SVZs shows that the decrease in quiescent NSCs observed at 4 weeks post-*Cx43* deletion is not due to abnormal astrogliosis (**Extended Figure 3a**). We also observed significantly increased EGFR^+^SOX2^+^EdU^+^ activated NSCs in both mutant SVZs at 1-week post-Tx (**Figure 3d and 3j**); however, at 4 weeks post-Tx, the number of activated NSCs was significantly decreased (**Figure 3g and 3j**). Neuroblasts, identified via expression of DCX, were significantly increased in both *Cx43*EC^iKO^ and *Cx43*Glast^iKO^ SVZ at 1-week post-*Cx43* deletion (**Figure 3e and 3k**), while their number was significantly reduced in both mutants at 4 weeks post-Tx (**Figure 3h and 3k**). Furthermore, we used *Cx43*Glast^iKO^;ROSA^mT/mG^ mice to perform lineage-tracing at 1 week post-*Cx43* deletion. We found DCX^+^ neuroblasts (**Extended Figure 3b**, white arrow heads) localized within the recombinant GFP^+^ population, supporting that the neuroblasts generated in the *Cx43*Glast^iKO^ SVZ (**Figure 3e and 3k**) were derived from *Cx43-*deficient NSCs.

Elsewhere in the brain, with exception of the SVZ, vascular ECs interact with astrocytes and pericytes via Cx43, which promotes their blood-brain-barrier (BBB) function ^36–38^. Thus, we sought to determine whether deletion of *Cx43* in vascular ECs or Glast-expressing astroglial cells compromised BBB integrity and caused vascular leakage. To assess vascular permeability, we injected *Cx43^fl/fl^*, *Cx43*EC^iKO^ and *Cx43*Glast^iKO^ mice with 2% Evans blue 24 hr prior to sacrifice, as depicted in **Extended Figure 4a**, and evaluated vascular leakage in brain and liver tissues. As expected, we did not observe vascular leakage in the control brains; there was also no leakage in the mutant brains (**Extended Figure 4b**). In contrast, in liver tissues that lack barrier function, leakage of Evans blue dye from the vasculature was evident throughout control and mutant tissues (**Extended Figure 4c**). In addition, using Vascupaint green perfusion, we did not observe any differences in *Cx43*EC^iKO^ and *Cx43*Glast^iKO^ brain microvasculature morphology at 4 weeks post-Tx, compared to *Cx43^fl/fl^* controls. Thus, deletion of *Cx43* in vascular ECs or astroglial cells does not appear to compromise BBB integrity in the brain. We also measured brain and OB areas and found no differences between *Cx43^fl/fl^* control mice and *Cx43*EC^iKO^ and *Cx43*Glast^iKO^ mice after short- or long-term deletion of *Cx43* **(Extended Figure 5).**

### Cx43 deletion in ECs or NSCs increases neuroblast generation in the RMS and neurogenesis in the OB

Since neuroblasts generated in the SVZ migrate toward the OB via the RMS, we next assessed whether increased neuroblast generation in the *Cx43*EC^iKO^ and *Cx43*Glast^iKO^ SVZs at 1-week post-Tx leads to increased neuroblasts in the RMS (**Figure 4a**). On brain sagittal sections immunostained with anti-DCX, we analyzed the anterior portion of the RMS recognized by its elbow-shaped morphology (**Figure 4b and 4c**). In this region, we found significantly increased DCX^+^ neuroblasts in *Cx43*EC^iKO^ and *Cx43*Glast^iKO^ mice, compared to *Cx43^fl/fl^* controls (**Figure 4d and 4e**). We next examined whether the observed increase in neuroblasts in *Cx43*EC^iKO^ and *Cx43*Glast^iKO^ RMS at 1-week post-Tx affects neurogenesis in the OB. To do so, we used a LRC protocol (**Figure 4f**), in which 6-week-old adult mice received 3 consecutive EdU injections on the first 3 days of a 5-day Tx-induction. At 4 weeks post-Tx, mice were analyzed via immunostaining with anti-EdU and anti-NeuN, to label newborn neurons^2, 39^. Interestingly, we observed a significantly increased number of LRC/NeuN^+^ interneurons in the granule cell layer (GCL) of both *Cx43*EC^iKO^ and *Cx43*Glast^iKO^ OB compared to *Cx43^fl/fl^* controls (**Figure 4g-4i**).

**Figure 4:**
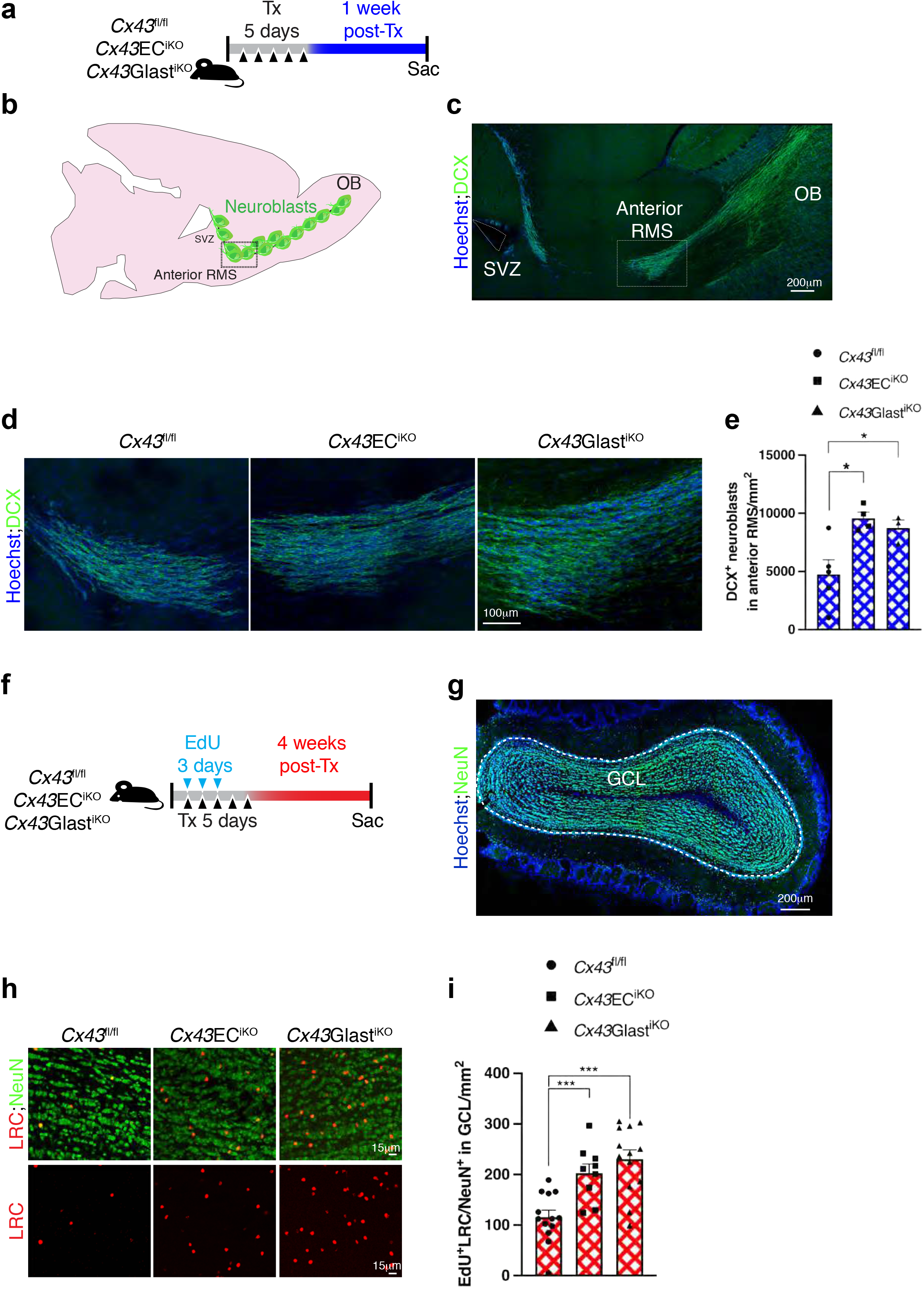
Cx43 deletion in ECs or NSCs increases neuroblast generation in the RMS and neurogenesis in the OB. (**a**) Timeline used to evaluate the effects of short-term deletion of *Cx43* on neuroblasts in the RMS. (**b**) Schematic representation of a mouse brain coronal section (50 μm) showing the RMS. (**c**) a representative confocal image of DCX^+^ neuroblasts in the RMS. Dashed boxes in (**b**) and (**c**) highlight the anterior RMS. (**d**) Representative confocal images of DCX^+^ neuroblasts in the anterior RMS of *Cx43*^fl/fl^, *Cx43*EC^iKO^ and *Cx43*Glast^iKO^ mice. (**e**) Quantifications of images shown in (**d**) (n=4 different images analyzed from 2 animals per group). (**f**) Timeline used to evaluate the effects of long-term deletion of *Cx43* on neurogenesis in the olfactory bulb. (**g**) Representative confocal image of NeuN^+^ newborn neurons (in green) on a coronal section (50 μm) of the mouse olfactory bulb. The dashed white lines outline the granule cell layer (GCL). (**h**) Representative images of EdU^+^ label retaining cells (LRC, red) and NeuN^+^ newborn neurons (green) in GCL of *Cx43*^fl/fl^, *Cx43*EC^iKO^ and *Cx43*Glast^iKO^ mice. (**i**) Quantifications of images shown in (**h**) (n=5-8 sections analyzed from 2 different animals per group). Scale bars are 200 μm in (**c**), 100 μm in (**d**) and (**g**) and 15 μm in (**h**). Data are mean ± SEM. \**p* ≤ 0.05, \*\*\**p* ≤ 0.001.

To determine whether neurons in the *Cx43*Glast^iKO^ RMS were derived from SVZ NSCs, we used *Cx43*Glast^iKO^;*ROSA^mT/mG^* mice to perform lineage-tracing studies (**Extended Figure 6a**). We found LRC/NeuN^+^ cells (**Extended Figure 6b**, white arrows) localized within the recombinant GFP^+^ population, supporting that the newborn olfactory neurons generated in the *Cx43*Glast^iKO^ SVZ (**Figure 3e and 3k**) were derived from *Cx43-*deficient NSCs. Importantly, the increase in newborn neurons in *Cx43*EC^iKO^ and *Cx43*Glast^iKO^ OB was not a consequence of neuronal death, as we did not observe changes in apoptosis levels in *Cx43*EC^iKO^ and *Cx43*Glast^iKO^ OB measured via immuno-staining for cleaved CASPASE-3 (**Extended Figure 6c-e**). Collectively, these results suggest that the short-term deletion of *Cx43* in ECs or NSCs leads to decreased quiescent NSCs and increased activated NSCs in the SVZ, as well as increased neuroblast generation in the SVZ and RMS. After long-term deletion of EC- or NSC-expressed *Cx43*, the quiescent NSC pool in the SVZ is further depleted, while activated NSCs and neuroblasts are exhausted from the SVZ, and neurogenesis is increased in the OB. Thus, both EC- and NSC-expressed Cx43 contributes to the regulation of adult SVZ neurogenesis.

### EC-expressed Cx43 is necessary for SVZ niche repopulation

Our studies suggested that NSC quiescence and maintenance in the SVZ niche is regulated by ECs in a Cx43-dependent manner. Thus, we tested whether lack of EC-expressed Cx43 would impair NSC-mediated SVZ niche repopulation after depletion of all SVZ proliferating cells (activated NSCs, TACs and neuroblasts) via anti-mitotic drug Ara-C (cytosine-β−D-arabinoside). At 1-week post-Tx injections in *Cx43*EC^iKO^ and *Cx43^fl/fl^* control mice, Ara-C (2%) or saline (control) was infused directly in one of the two lateral ventricles (ipsilateral) via a cannula connected to a subcutaneously implanted mini-osmotic pump for 6 consecutive days (**Figure 5a**). EdU was administered to label actively proliferating cells 24 hr prior to sacrifice. The intracerebroventricular injection coordinates were verified via injection of 1% Fast Green dye, as previously described^40^ (**Extended Figure 7a**).

**Figure 5:**
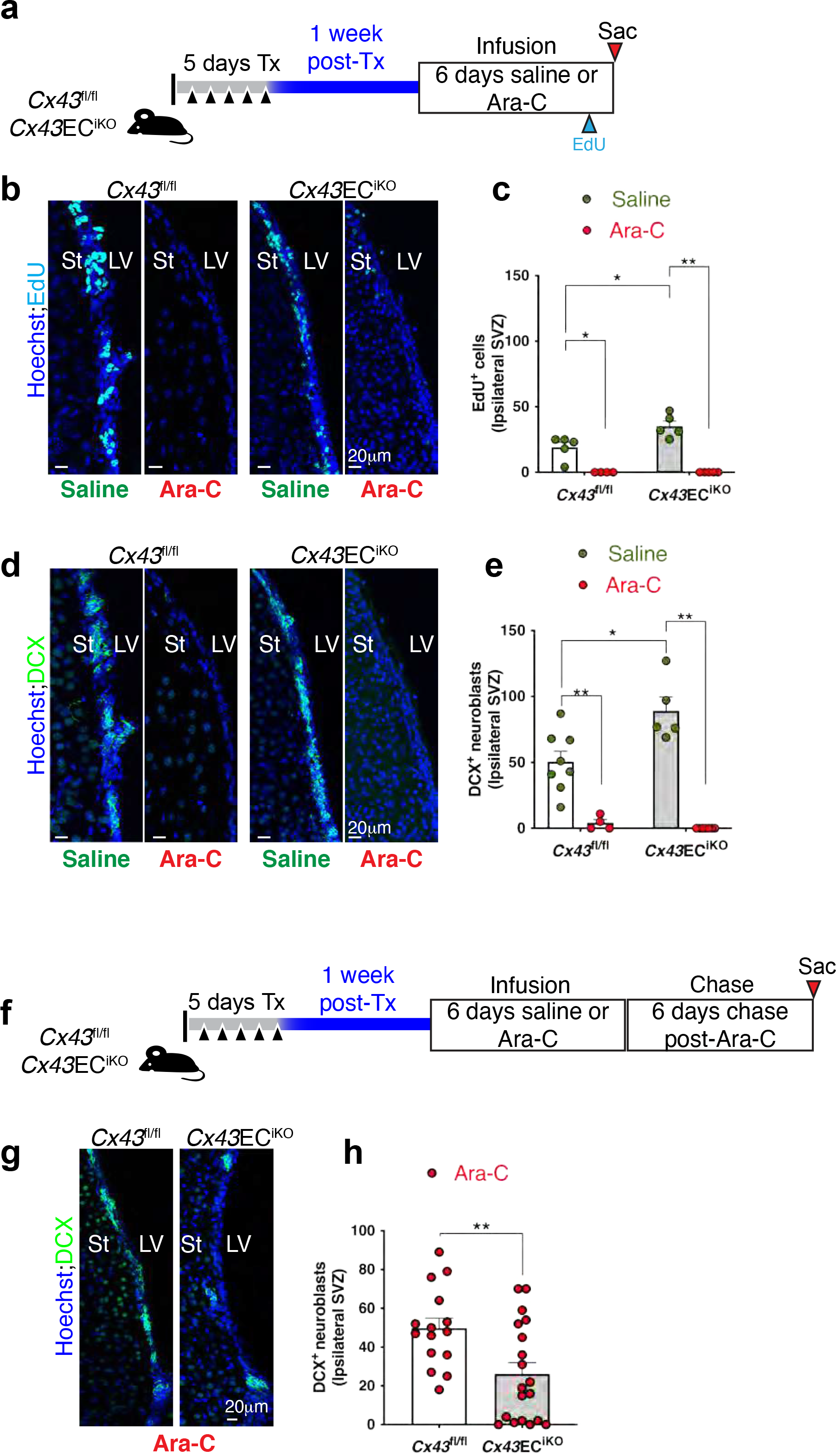
EC-expressed Cx43 is necessary for SVZ niche repopulation. (a) Timeline used for saline or Ara-C infusions. (**b**) Representative confocal images showing EdU^+^ cells in SVZ of saline vs. Ara-C infused *Cx43*^fl/fl^ and *Cx43*EC^iKO^ mice after 6 days infusion. (**c**) Quantifications of images shown in (**b**) (n=2 different animals). (**d**) Representative confocal images showing DCX^+^ neuroblasts in SVZ of saline vs. Ara-C infused *Cx43*^fl/fl^ and *Cx43*EC^iKO^ mice. (**e**) Quantifications of images shown in (**d**) (n=2 biological replicates). (**f**) Timeline used for Ara-C infusion and chase period. (**g**) Representative confocal images showing DCX^+^ neuroblasts in SVZ of saline vs. Ara-C infused *Cx43*^fl/fl^ and *Cx43*EC^iKO^ mice after Ara-C infusion and chase period. (**h**) Quantifications of images shown in (g) (n=3 biological replicates). Scale bar is 20 μm. St: Striatum; LV: Lateral ventricle. Data are mean ± SEM. \**p* ≤ 0.05, \*\**p* ≤ 0.01.

We first confirmed that Ara-C treatment depleted all EdU^+^ proliferating cells (**Figure 5b and 5c**), as well as DCX^+^ neuroblasts (**Figure 5d and 5e**), in the SVZ. Importantly, we observed increased DCX^+^ neuroblasts in control, saline-infused *Cx43*EC^iKO^ SVZ compared to *Cx43*^fl/fl^ (**Figure 5d and 5e**). These results are consistent with our previous studies (**Figure 3e and 3k**) where we demonstrated that short-term deletion of *Cx43* in *Cx43*EC^iKO^ leads to increased SVZ neuroblasts. Subsequently, we analyzed the number of DCX^+^ neuroblasts in the SVZ 6 days after Ara-C withdrawal (**Figure 5f**, chase period). DCX^+^ neuroblasts were significantly reduced in *Cx4*3EC^iKO^ SVZ, compared to *Cx43*^fl/fl^ (**Figure 5g and 5h**). Thus, absence of *Cx43* in ECs impairs the re-population of the SVZ niche post-Ara-C treatment, supporting that Cx43 expression in ECs contributes to the maintenance of the quiescent NSCs pool in the SVZ and prevents their premature activation and depletion.

### Cx43 cytoplasmic tail mediates EC-induced NSC quiescence in an ERK-dependent manner

We next investigated the mechanism(s) by which Cx43 regulates NSC-EC interactions to maintain NSC quiescence. Cx43 can function in a channel-dependent or channel-independent manner; thus, we created lentiviral constructs to express Cx43 mutant proteins to perform structure-function studies to determine how Cx43 mediates NSC-EC interactions. To investigate channel-dependent functions, we generated a Cx43 channel-dead mutant (*Cx43T154A*) that allows gap junction channel formation but blocks Cx43 channel activity^41^. To investigate channel-independent functions, we generated mutant Cx43 that lacks its cytoplasmic tail (*Cx43CTΔ258*)^42^, which can mediate intracellular signaling, independent of channel formation^43^. ECs and NSCs were treated with si*Cx43* to suppress endogenous Cx43 expression, and then transduced with lentiviral constructs to express the mutant Cx43 proteins. Expression of the Cx43 cytoplasmic tail truncated mutant (*Cx43CTΔ258*, 1 μg), lead to a significant downregulation of genes associated with NSC quiescence (*Gfap, Nestin* and *Glast*), while expression of the Cx43 dead-channel mutant (*Cx43T154A*, 0.5 μg) had no effect on NSC gene expression (**Figure 6a, Extended Figure 8a and 8b**). Additionally, when we treated both NSC and EC with ^43^gap 26 peptide (100 nM), which blocks Cx43 channel activity, we did not observe any changes in NSC gene expression associated with either quiescence or activation (**Figure 6b**). Thus, it appears that EC-mediated NSC quiescence is regulated by Cx43 in a channel-independent manner. We assessed whether overexpression of *Cx43* in NSCs alone was sufficient to suppress NSC proliferation, as measured via EdU incorporation, and found that Cx43 overexpression in NSC does not affect their proliferation (**Extended Figure 8c-g**).

**Figure 6:**
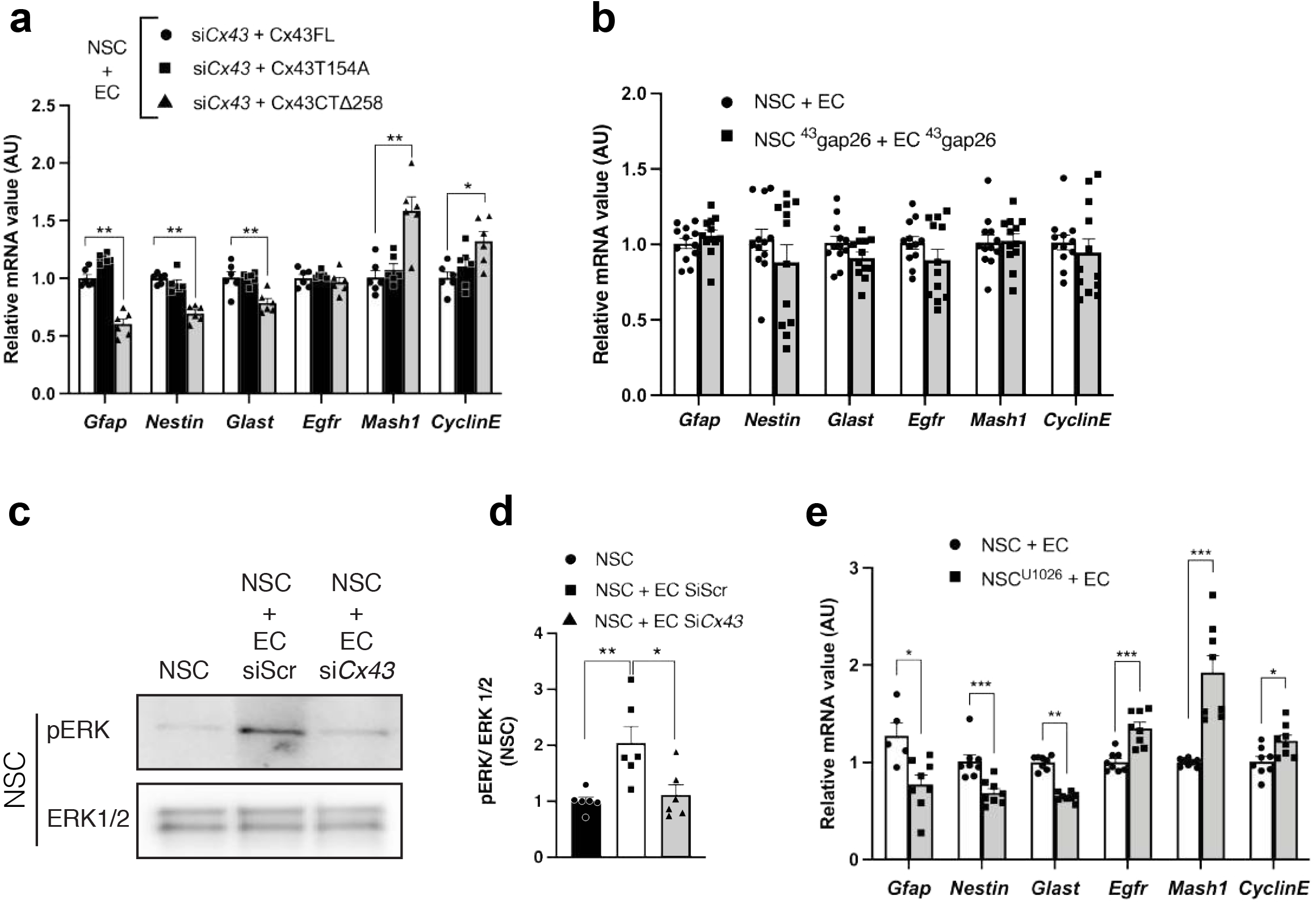
Cx43 cytoplasmic tail mediates EC-induced NSC quiescence in an ERK-dependent manner. (**a**) qPCR analysis of quiescent and activated genes in NSCs co-cultured with ECs where both cell types are treated with Si*Cx43* followed by either the *Cx43*T154A mutant (Cx43 dead channel) or the *Cx43*CTΔ258 mutant (cytoplasmic tail truncated) compared to NSCs, and ECs co-culture treated with Si*Cx43* followed by *Cx43* full length (FL) construct. (n=3 different experiments). (**b**) qPCR analysis of quiescent and activated NSCs co-cultured with ECs where both cell types are treated with ^43^gap26 compared to NSCs and ECs co-culture (n=3 different experiments). (**c**) Western blot analysis of pERK and ERK protein levels in NSCs cultured alone, or co-cultured with ECs treated with control siRNA (SiScr) or *Cx43* siRNA (Si*Cx43*) and quantifications are shown in (**d**) (n=6 different experiments). (**e**) qPCR analysis of quiescent and activated NSCs co-cultured with ECs where NSCs is treated with the ERK signaling inhibitor U0126 compared to NSCs and ECs co-culture (n=2 different experiments). Data are mean ± SEM. \**p* ≤ 0.05, \*\**p* ≤ 0.01, *** *p* ≤ 0.001. **** *p* ≤ 0.0001.

**Figure 7:**
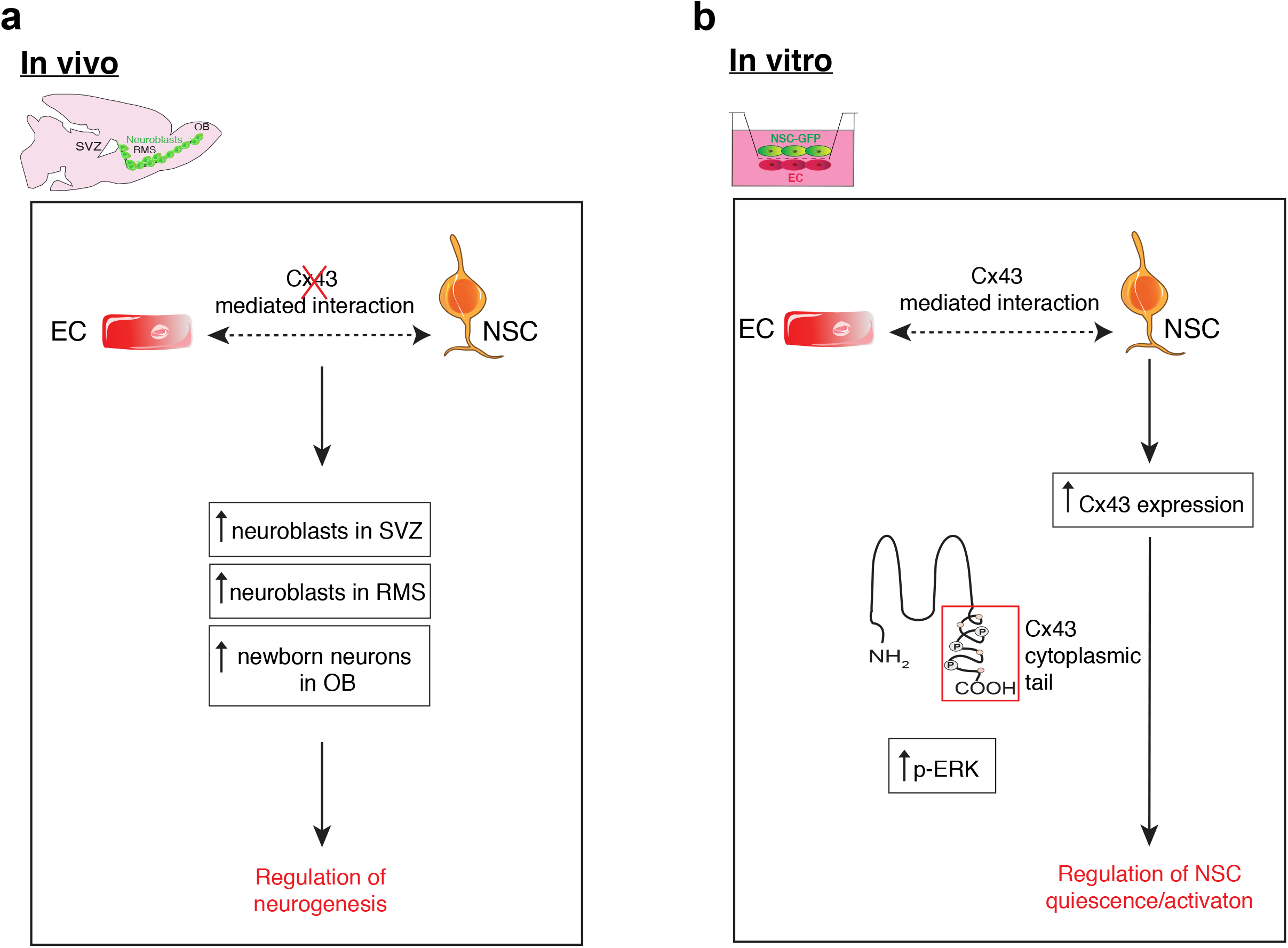
Schematic representation of Cx43 regulation of NSCs. (a) in vitro: ECs and NSCs co-culture upregulates Cx43 expression. Cx43-mediated regulation of NSC quiescence and activation is dependent on Cx43 cytoplasmic tail and ERK activation. (**b**) in vivo: absence of Cx43-mediated interaction between ECs and NSCs increases the number of neuroblasts in the SVZ and RMS leading to an increase of newborn neurons in the OB. Absence of vascular Cx43 impairs SVZ neuroblasts re-population post depletion.

It is known that the Cx43 cytoplasmic tail is involved in the activation of downstream effectors involved in ERK signaling^43, 44^ and, interestingly, we observed increased ERK activation in NSCs when co-cultured with ECs and the latter is abrogated in NSCs co-cultured with ECs silenced for *Cx43* (**Figure 6c and 6d**). Finally, inhibition of ERK signaling with U0126 (10 μM) in NSCs co-cultured with ECs abolished the effect of ECs on NSC expression of genes associated with quiescence (**Figure 6e**). Collectively, these results suggest that EC maintenance of NSC quiescence is mediated by the Cx43 cytoplasmic tail, via ERK activation in NSCs.

## Discussion

NSCs in the adult brain SVZ reside in a vascularized niche, which is known to regulate their proliferation, migration, and differentiation via niche cell-derived secreted factors^21, 22, 45–49^. It is also well established that quiescent NSC basal end-feet are in direct contact with vascular ECs allowing neuro-vascular interactions^7^. Past studies showed that the expression of Cx43 in neural progenitor cells maintains their survival and proliferative state^27, 50–53^. However, the role of Cx43-mediated neuro-vascular interactions in the adult SVZ in the regulation of NSC behavior has not been investigated.

In this study, we first used an in vitro EC-NSC co-culture system that allows direct cell-to-cell contact^54, 55^ to demonstrate that ECs decrease NSC proliferation and increase their expression of genes associated with quiescence in a Cx43-dependent manner. We then used transgenic mice to show that deletion of Cx43 in ECs or NSCs in vivo leads to increased NSC proliferation and neuroblast generation in the SVZ, as well as increased neurogenesis in the OB. Our observations are consistent with previously published studies showing that the vascular niche maintains quiescent NSCs and promotes their survival^25, 56^, and provide insight into the underlying mechanisms of regulation.

Interestingly, our in vivo results show the same outcome whether Cx43 is deleted in ECs or NSCs while, in vitro, only EC-expressed Cx43 mediates effects on NSC proliferation and survival. There are many differences between the two systems that can account for this finding, including lack of other SVZ components in NSC-EC co-cultures that may be important for this regulation. In addition, in the in vitro studies, NSCs were silenced for *Cx43* prior to co-culture with ECs while in vivo NSCs and ECs were in contact prior to *Cx43* deletion in either ECs or NSCs. Thus, there are likely more complex interactions between ECs and NSCs, and perhaps other cell types, in the SVZ niche that depend on EC- and NSC-expressed Cx43.

The evaluation of the short- vs. long-term deletion of *Cx43* in either ECs or NSCs showed that the SVZ quiescent NSC pool is gradually depleted over time and leads to increased neuroblasts in the RMS and ultimately increased newborn neurons in the OB. Consistent with this, when quiescent NSCs in the *Cx43*EC^iKO^ SVZ were triggered towards activation to replenish the depleted niche post-AraC infusion, the number of neuroblasts generated in *Cx43*EC^iKO^ was significantly less than controls. This finding is consistent with the idea that deletion of *Cx43* in ECs activates quiescent NSCs in the adult SVZ which leads to a gradual exhaustion of the NSC pool over time; thus, the NSCs in the *Cx43*EC^iKO^ SVZ cannot replenish the niche to the same extent as control littermates. Whether the increased newborn neurons in the OB leads to functional neurons that fully integrate the olfactory neuronal circuits and modify the olfaction behavior of these Cx43- deficient animals is not yet known.

Using mutant Cx43 proteins, we demonstrated that Cx43 mediates EC regulation of NSCs in a channel-independent manner that involves its cytoplasmic tail and ERK activation. However, further mechanistic studies are needed to determine how Cx43 is regulating ERK activation in NSCs co-cultured with ECs. For example, whether the phosphorylation of Cx43 on a particular serine residue of the cytoplasmic tail is responsible for ERK activation remains to be addressed; however, the phosphorylation serine sites 255, 279 and/or 282 of the Cx43 cytoplasmic region were previously linked with ERK activity^57^. Also, it has been shown that activated ERK can bind to Cx43, suggesting that Cx43 may play a direct role in ERK regulation^58^. We found that EC-NSC co-culture increases Cx43 expression in NSCs; however, over-expression of Cx43 in NSCs does not mimic the effects of EC-expressed Cx43 on NSC behavior or gene expression. Thus, although it is possible that, in NSCs, ERK signaling via the Cx43 cytoplasmic tail enables the activation of downstream effectors that promote NSC quiescence, the regulatory mechanisms are likely to be more complex.

In summary, we used both in vitro and in vivo approaches to gain insight into the role of Cx43 in the regulation of NSCs in the adult SVZ. We show that EC- and NSC-expressed Cx43 are required to maintain NSC quiescence. Further mechanistic studies are needed to determine exactly how Cx43 is regulating NSC behavior in the adult SVZ, and our in vitro studies suggest that this occurs in a channel-independent manner. Such insights can be applied to the bioengineering of a NSC niche ex vivo that better mimics its in vivo environment, to enable sustained NSC viability and functional properties for stem cell therapies for neurological disorders.

## Acknowledgements

The authors of this work were supported by: NIH grants to K.K.H. (R01 EB 016629 and HL146056, and NIH U2EB017103), AHA post-doctoral fellowship to N.G. (19POST34400065), NIH grants to C.N. (T32 HL007224, T32 HL007284).

## Author contributions

N.G, G.G., J.S.F., N.C.W., H.H.V., J.S.G., B.R.A., N.B., K.B., G.M., performed experiments. S.M. and M.R.H. maintained mouse lines and performed mouse genotyping. B.R.A generated Cx43 lentiviral constructs. N.G., G.G. and K.K.H designed experiments, analyzed data and wrote the manuscript. A.E., J.L.T., C.F.C contributed to experimental design and manuscript editing.

## Material and Methods

### Animals

*Cdh5*Cre^ERT2^1Rha (also referred to as *VE-Cadherin*Cre^ERT2^)^59, 60^ and *GLAST*Cre^ERT2^ (*Slc1a3*Cre^ERT2^)^61^ were gifts from Drs. Ralf Adams and Jean Léon Thomas labs, respectively. *Gja1^flox/flox^* (also referred to as *Cx43^flox/flox^*) and *ROSA^mT/mG^* mice were commercially purchased. For genetic loss-of-function studies, *Cdh5*Cre^ERT2^ and *GLAST*Cre^ERT2^ animals were crossed to mice carrying a loxP-flanked *Gja1* gene (*Cx43^flox/flox^*) to create *Cx43*EC^iKO^ and *Cx43*Glast^iKO^, respectively. Mice were maintained under standard pathogen-free conditions. All animal protocols and procedures were reviewed and approved by the University of Virginia Animal Care and Use Committee (protocol #4277) and complied with all ethical regulations. To induce Cre activity in *Cx43*EC^iKO^ and *Cx43*Glast^iKO^, 6-week-old adult mice received intraperitoneal (i.p.) injections of Tamoxifen (Tx, 2mg/day) for 5 consecutive days, which resulted in ∼90% and ∼75% *Gja1* deletion in ECs of *Cx43*EC^iKO^ and NSCs of *Cx43*Glast^iKO^ respectively, as assessed by qPCR (Figure 3E). Tx-injected *Cx43^flox/flox^* littermates were used as controls. Mice of both sexes were used to minimize gender-related biased results and analyzed at 1 week-post final Tx injections for short-term deletion studies and 4 week-post final Tx injections for long-term deletion studies. For Ara-C studies, mice were analyzed at 2- and 3-week-post final Tx injection. To label actively proliferating progenitors in the SVZ, mice received an i.p. injection of EdU (50mg/kg) 24 hr prior to sacrifice. We used label retention to identify quiescent NSCs that are slow-cycling cells, also known as label retaining cells (LRC) that retain BrdU or EdU for extended periods due to their relatively long cycling times ^18, 62, 63^. To do so, mice received 5 consecutive injections of BrdU or 3 consecutive EdU injections on the first 3 days of the 5 days of Tx injections, mice were analyzed 4-weeks post-Tx.

### Mouse genotyping

Ear sample DNA was lysed using the hotshot method. Samples were lysed in alkaline lysis reagent (pH 12) containing 25mM NaOH and 0.2mM EDTA for 1 hr at 90 °C. Subsequently, samples were neutralized using 40mM Tris-HCl. PCR was performed in PCR-ready tubes (Bioneer Inc. K-2016) containing 1μL of DNA sample and 12.5 mM primers for a final reaction volume of 20μL adjusted with RNAse-free H_2_O. PCR genotyping primers sequences are documented in **Table 1**.

**Table 1:**
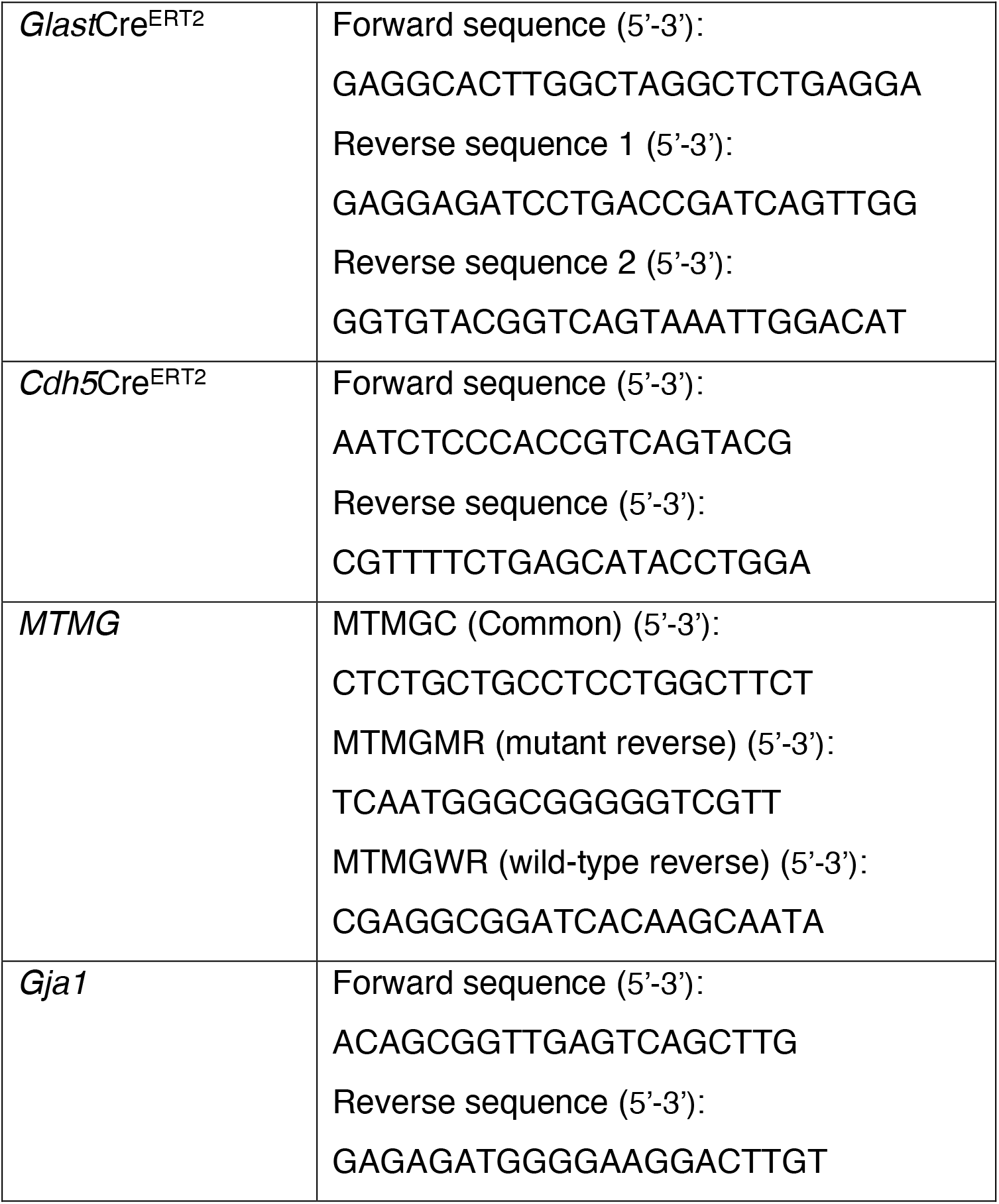
PCR genotyping primers sequences.

### Tissue collection, subventricular zone (SVZ), rostral migratory stream (RMS) and OB immunohistochemistry and imaging

Mice were sacrificed with a lethal dose of ketamine (80mg/kg body weight)/xylazine (8mg/kg body weight). Mice received trans-cardiac infusion of 10mL sterile PBS supplemented with 2mM EDTA and 10U/mL heparin followed by 10mL 3.7% formaldehyde. Brain tissues were post-fixed with 3.7% formaldehyde at 4°C overnight. Post-fixation, brains were washed 3 times during 10 min in PBS 1X. SVZ coronal sections, sagittal sections of the RMS and coronal sections of the OB were collected in a 24-well plate in a sequential manner and subsequently subjected to a free-floating slice immunostaining protocol. All sections were cut at 50 μm with a Leica vibratome (VT1000S). Briefly, slices were permeabilized with 0.5% Triton X-100 for 30 min at room temperature (RT). If EdU staining was required, we proceeded with EdU staining post-permeabilization following the manufacturer’s protocol (Click-iT™ – ThermoFischer C10337). Sections were then blocked with 10% donkey serum in PBS 1X supplemented with 0.3% Triton X-100 (blocking buffer) for 1 hr at RT. After blocking, sections were incubated with appropriate primary antibodies (listed in **Table 2**) diluted in blocking buffer overnight at 4°C, on a low-speed rocking plate. Samples were washed three times with PBS 1X supplemented with 0.1% Triton X-100 (PBST) then incubated with respective conjugated secondary antibodies for 1 hr at RT. After 3 washes with PBST, Hoechst (4 μM) was added during 30 min for nucleus counterstaining. SVZ images were acquired with an inverted Leica SP8 DMi8 high-resolution confocal microscope equipped with adaptive deconvolution (LIGHTNING^®^, Leica) using 63x or 20x objectives. A series of 3 to 5 SVZ slices from each animal taken from the same rostro-caudal area, judged by the shape of the lateral ventricles, of the corpus callosum and the anterior commissure were imaged. We localized the RMS attached to the OB between a sagittal depth of 3.2-3.35mm from the lateral side of the brain toward the midline. For OB analysis, the whole OB was cut coronally. Images were post-analyzed using ImageJ and Photoshop (Adobe) software.

**Table 2:** Primary antibodies used for SVZ immunostaining.

| Antibody/catalogue number | Species | Dilution |
| --- | --- | --- |
| Connexin 43 (Abcam ab11370) | Rabbit | 1:1000 |
| SOX2 (ebioscience 14-9811-82) | Rat | 1:100 |
| EGFR (Millipore 06-847) | Rabbit | 1:75 |
| Doublecortin (Abcam ab18723) | Rabbit | 1:200 |
| Doublecortin (AVES lab DCX) | Chicken | 1:200 |
| GFAP (DAKO Z0334) | Rabbit | 1:400 |
| GFAP (AVES lab GFAP) | Chicken | 1:400 |
| PECAM1 (CD31) (Millipore MAB1398Z) | Hamster | 1:100 |
| NeuN (Abcam ab104225) | Rabbit | 1:500 |
| S100β (DAKO Z0311) | Rabbit | 1:300 |
| CASPASE-3 (Cell signaling 9661S) | Rabbit | 1:200 |

### Evans blue permeability assay

To assess changes in vascular permeability, mice received an i.p. injection of 2% Evans blue and were sacrificed 24 hr later. Prior to brain and liver harvesting, mice were infused with PBS 1X supplemented with 2mM EDTA and 10U/mL heparin to wash out any traces of blood and maintain vessel integrity. Brains were post-fixed in 3.7% formaldehyde overnight and imaged using a dissecting microscope equipped with a digital camera (Leica).

### Vacupaint™ Silicone Rubber injection

To assess changes in the brain microvaculature, mice were injected with 4 ml of Vascupaint (green) following the manufacturer’s protocol (ediLumine™). Post-vascupaint injection, brains were harvested and post-fixed in 3.7% formaldehyde overnight. Brains were then dehydrated with 30%, 60% and 100% methanol successively for 24h each. Post-dehydration, brains were clarified with 50:50 of Benzyl Benzoate:Benzyl alcohol for 48h and imaged using a Nikon SMZ-745T Trinocular 4K Digital Stereo microscope.

### Ara-C infusions

Cytosine-β-D-arabinofuranoside (Ara-C) infusions were performed as previously published^6, 64^. Briefly, 2% Ara-C (Sigma) in 0.9% saline or saline alone was infused directly in the lateral ventricle (ipsilateral side) of adult mice (8-week-old) via a cannula (Alzet^®^ brain infusion kit 3) implanted stereotaxically in a 1mm burr hole drilled on the surface of the brain at the following coordinates: 1.4 mm lateral and 0.5 mm rostral to bregma. Intracerebroventricular injection coordinates were verified with the injection of Fast Green dye 1% as previously described^40^ (**Extended Figure 5**). The cannula was connected to a subcutaneously implanted mini-osmotic pump (Alzet^®^ model 1007D flow rate 0.5μl/hr, 7 days). After 6 days of infusion, the pump was removed, and mice were sacrificed at the indicated survivals. The success of the Ara-C infusion was evaluated by immunostaining for the neuroblast marker, DCX and the proliferation marker EdU.

### FACS isolation of primary ECs and NSCs from mouse brain cortex and SVZ

Primary ECs and NSCs from mouse brain cortex and SVZ were FACS-isolated following a detailed published protocol^65^. Briefly, brain cortex and SVZ were harvested (5 brains at a time to maximize cell viability), minced into small pieces of ∼1 x 1 mm in size. Minced pieces are then subjected to enzymatic dissociation in collagenase/dispase (Roche, 100mg/ml stock) solution for 30 min at 37°C in a hybridization oven with constant rotation. Subsequently, the digested tissues were triturated in 2% FBS in PBS supplemented with DNAse (10 mg/ml stock solution) with a P1000 pipette (∼100x). Afterward, debris and myelin were removed with Percoll, cells were pelleted by centrifugation and resuspended in HBSS/BSA/glucose buffer for immunostaining. Antibodies (**Table 3**) were added to the resuspended cells (1 μl of antibody/10^5^ cells) and incubated during 20 min on ice protected from light. Post-immunostaining cells were washed with HBSS/BSA/glucose buffer and filtered through a 40 μm cell strainer. Cells isolated from the cortex were used as unstained, single-color and FMO controls (each combination of all antibodies except one), as well as isotype controls to set up gating and compensation strategy (**Extended Figure 3**). From SVZ CD45^-^ cells, ECs (CD31^+^Glast^-^ population) and NSCs (Glast^+^CD31^-^ population) were collected using a BD FACS Melody cell sorter equipped with a 100 μm nozzle in pre-filled FACS tubes with EC and NSC media respectively to cushion the cells. Post-FACS processing, cells are pelleted, snap frozen in liquid nitrogen and stored at -80 °C for RNA isolation.

**Table 3:**
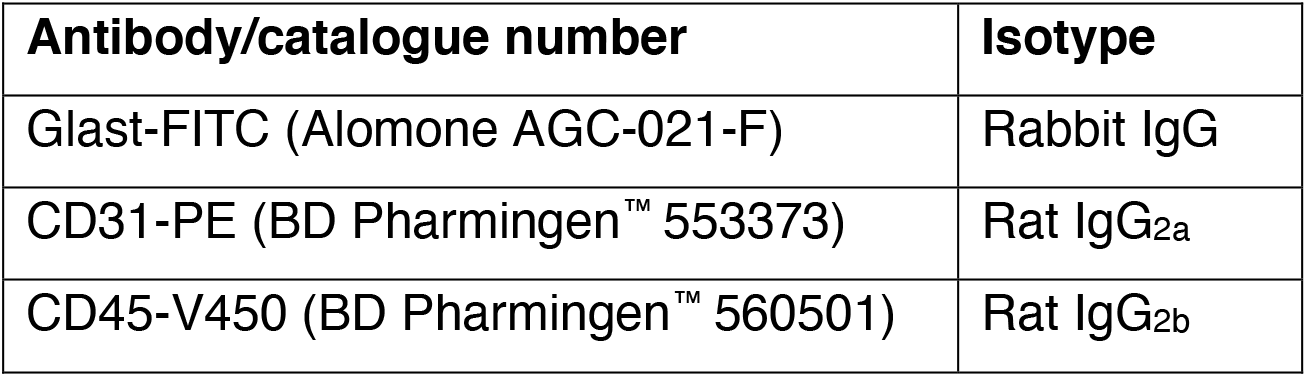
Antibodies used for FACS.

### RNA isolation

RNA from primary ECs and NSCs, as well as bEnd.3 and ANS4-GFP cells, were purified using RNeasy Plus micro kit (Qiagen). 1 μg of RNA was reverse transcribed using high-capacity cDNA Reverse transcription kit (Applied Biosystems™). Quantitative PCR was performed on 15 ng cDNA using PowerUp SYBR Green Master Mix (Applied Biosystems™) and the corresponding primers (**Table 4**). The data were first normalized to actin level in each sample, and the relative expression levels of different genes were calculated by the comparative Ct method^66^.

**Table 4:**
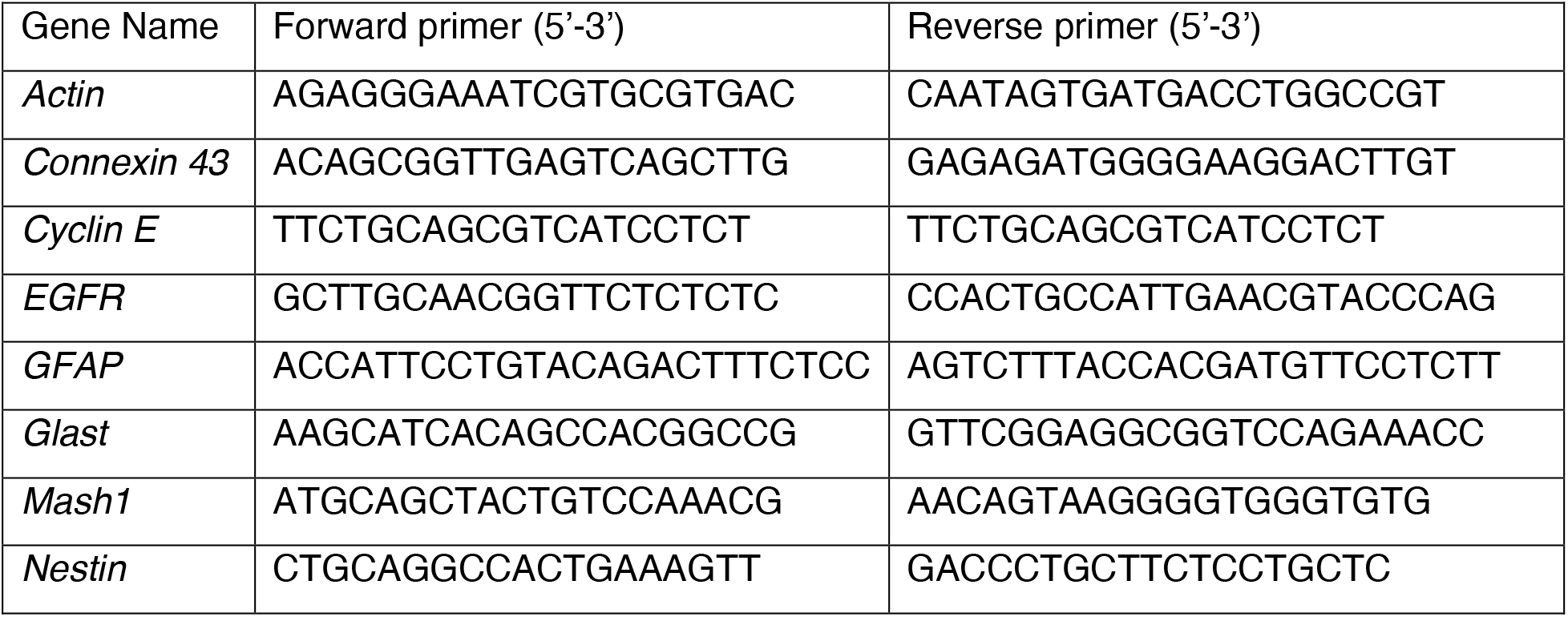
Mouse primers used for quantitative PCR.

### siRNA transfection

*Cx43* siRNA (Smartpool siRNA *Cx43*, L-051694-00-0005) and the negative control/SiScramble (Non-targeting pool siRNA, D-001810-10-05) were purchased from Dharmacon. We transfected bEnd.3 cells when 70% confluent with 60nM siRNA per six-well plate. Experimentally, lipofectamine RNAimax (Invitrogen) is mixed with opti-MEM media (Gibco) and incubated at room temperature for 5 min (mix A). 60nM of Si*Cx43* premixed Opti-MEM is then added to mix A and incubated for 15 min (mix B). Mix B is then added to bEnd.3 cells DMEM media (ATCC) without penicillin/streptomycin. The same protocol was used to transfect ANS4-GFP cells with 40nM of Si*Cx43*. 60nM and 40nM of SiScramble (control) were used to transfect bEnd.3 and ANS4-GFP cells, respectively, for control experiments. Cells were used for experiments 72 hr post-transfection.

### *Gja1* overexpression in ANS4-GFP

*Gja1* overexpression in ANS4-GFP was performed using a pcDNA3.1^+^ /C-(K)DYK vector containing *mus musculus Gja1* (*Cx43*) cDNA (GenScript NM_010288.3). Transfection was performed according to the manufacturer’s experimental protocol in 6-well plate. To assess proliferation, EdU (5μM) was added to the culture media 24 hr post-transfection. At 48 hr post-transfection, NSC-GFP were collected for qPCR and western blot analysis to assess transgene expression or fixed for immunofluorescence studies to assess Cx43 expression and EdU uptake.

### Generation of Cx43 mutants

Human *GJA1* (gene for *Cx43*) full length (FL) construct, Cx43 cytoplasmic tail truncated at amino acid 258 (*Cx43ΔCT258*) and dead channel Cx43 mutant (*Cx43T154A*, mutation that converts threonine 154 into alanine) were PCR-amplified from pTRE-TIGHT-Cx43-eYFP (gifted from Robin Shaw; Addgene Plasmid #31807) and cloned into the pcDNA3.1-HA (gifted from Oskar Laur; Addgene Plasmid #128034). We inserted the PCR fragments at NheI/BamHI sites by infusion HD-cloning (Takara Biosciences), to excise HA-tag from the plasmid, rendering the constructs expressed as tag-less. Site-directed mutagenesis was achieved using Q5-Site-Directed Mutagenesis Kit (NEB, USA), according to the manufacturer’s protocol. Primers were designed using NEB-Base changer software (**Table 5**). Plasmids were sequenced and confirmed by Eurofin Sanger Sequencing Services, USA.

**Table 5:**
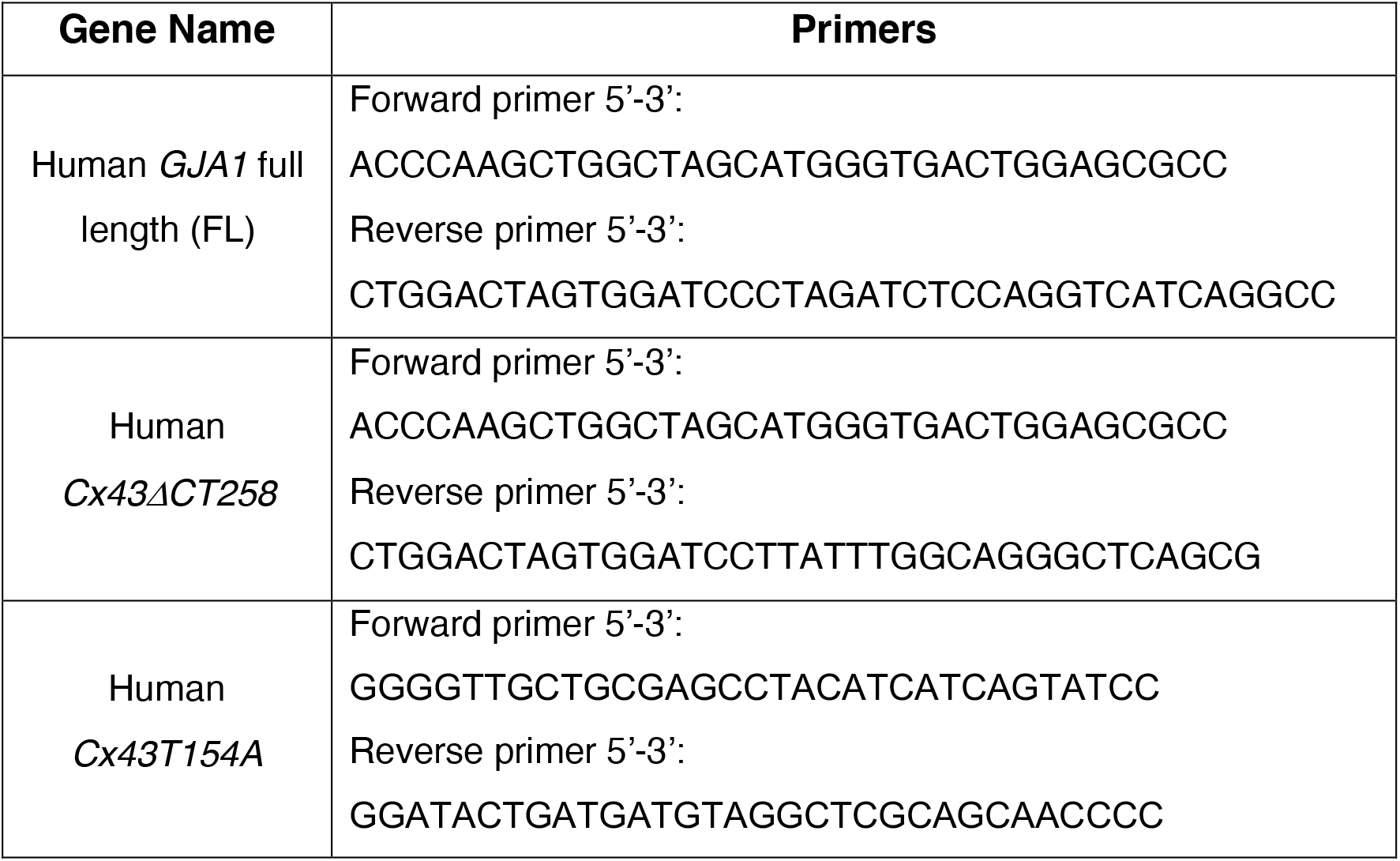
Human *Cx43* full length (FL), *Cx43ΔCT258* and *Cx43T154A* primers sequences.

### Cell culture

ANS4-GFP cells were gifted from Dr. Steve M. Pollard (University of Edinburgh). They were cultured as previously described^67–69^. bEnd.3 cells were purchased from (ATCC CRL-2299™) and cultured in Dulbecco’s Modified Eagle’s Medium (DMEM) (ATCC 30-2002) supplemented with 10% FBS and 1% penicillin/streptomycin. ANS4-GFP cells were used up to passage (P) 30 and bEnd.3 up to P16.

### Transwell co-culture of bEnd.3 and ANS4-GFP

Transwell polycarbonate membranes of a 12-well plate (Corning, catalogue #3401) were activated with media and bottom sides were pre-coated with 0.1% gelatin for 1 hr at 37°C. The transwell inserts were then placed with bottom side facing up in a 100mm tissue culture dish pre-filled with DPBS and bEnd.3 were then seeded. One hour later, inserts were flipped back in the wells of the 12-well plate pre-filled with bEnd.3 media and top side was coated with laminin (10μg/ml overnight at 37 °C). 24 hr-post-bEnd.3 seeding, the membrane was washed on both sides with serum free media to remove any traces of serum that may cause differentiation of ANS4-GFP cells. ANS4-GFP cells were then seeded on the top side of membrane in ANS4 media without growth factors (EGF and FGF). bEnd.3 and ANS4 were co-cultured at a 1:2 ratio in 100% ANS4-GFP media (supplemented with 5μM EdU when needed). Membranes designated for ANS4-GFP monocultures were subjected to the same extra-cellular matrix coating proteins as co-culture studies. ANS4-GFP were harvested after 48 hr of co-culture with trypsin for either qPCR analysis or fixed with 3.7% pre-warmed formaldehyde for immunofluoresence studies. For *Gja1* knock-down studies: bEnd.3 cells seeded on the bottom side of the membrane were transfected 24 hr before seeding ANS4-GFP cells on the top side and subjected to 48 hr of co-culture prior to harvest. For studies using si*Cx43* transfected ANS4-GFP: ANS4-GFP were transfected 24 hr prior to seeding on the top side of the membrane and harvested after 48 hr of co-culture. For studies using human *GJA1* mutants, ANS4-GFP and bEnd.3 were treated concomitantly with Si*Cx43* (see section siRNA transfection) and/or Cx43FL (1μg), Cx43T154A (0.5 μg), Cx43CTΔ258 (1 μg) for 6 hr. After Si*Cx43* and DNA transfection, cells were subjected to a media change and harvested after 48 hr of co-culture. To block Cx43 channel activity, after 24 hr of co-culture, ANS4-GFP and bEnd.3 cells were both treated with 100nM of ^43^gap 26 peptide (VCYDKSFPISHVR) (Genscript, catalogue # RP20274), every 8 hr for 24 hr. To block ERK signaling in the co-culture system, ANS4-GFP cells were treated with 10μM of U0126 (Cell signaling catalogue #9903S) for 24 hr. ANS4-GFP were collected (6-8 wells are pooled per condition) either for qPCR or Western blot analysis after 48 hr of co-culture.

### Immunofluorescence of transwell membranes

Insert wells were washed with DPBS supplemented with Ca^2+^ and Mg^2+^ allowing for cells to remain attached to the membrane. Cells were fixed with pre-warmed 3.7% formaldehyde in DPBS supplement with Ca^2+^ and Mg^2+^ for 15 min, then permeabilized with 0.5% Triton X-100 in PBS at room temperature for 20 min prior to EdU staining according to the manufacturer’s protocol (Click-iT™, Invitrogen). Cells were then blocked with 10% donkey serum and 0.1% Triton X-100 (blocking buffer) for 1 hr at RT followed by incubation with primary antibodies, chicken anti-GFP (1:500, Abcam ab13970) for NSCs and hamster anti-CD31 (1:500, Millipore MAB1398Z) for bEnd.3 cells, in blocking buffer at 4°C overnight and corresponding conjugated secondary antibodies for 1 hr at RT. Hoechst (5 μg/ml) was used for nuclear counterstaining. Membranes were cut out from inserts and mounted with DAKO mounting media (DAKO). Images were acquired with a DMi8 SP8 confocal microscope at 4 different viewing fields and analyzed in Image J.

### Bulk RNA sequencing

mRNA samples from ANS4-GFP and ANS4-GFP cells co-cultured with bEnd.3 (for 72 hours) were isolated using RNeasy Plus Micro Kit (QIAGEN, cat# 74034). Next-generation whole transcriptome Illumina sequencing (HiSeq4000) was performed by the Yale Center for Genome Analysis. Fastq-files of raw reads were generated with bcl2fastq2_v2.19.0 and then uploaded to the usegalaxy.org platform^70^ for quality control and trimming (Galaxy ToolShed v 1.0.2; FASTQ/A short-reads pre-processing tools: http://hannonlab.cshl.edu/fastx_toolkit/)^71, 72^. Sequences were aligned to the hg38genome using Kallisto ^73^ and aligned sequences were quantified with Sleuth ^74^. Differential gene expression analysis was performed with the Sleuth package in R by the Likelihood Ratio Test with regression of the experiment number to account for batch effects.

### Western Blot

Cells were lysed in RIPA buffer (Abcam, ab206996) and equal amounts of proteins (quantified according to the Pierce™ BCA™ assay kit manufacture’s protocol, ThermoFischer, 23252) were separated on 4-15% gradient Criterion precast gels (Bio-Rad 567-1084). Proteins were then transferred onto nitrocellulose membranes (Bio-Rad). Western Blots were developed with chemiluminescence HRP substrate (Radiance plus, Azure Biosystems AC2103) on a digital image analyzer, Azure Imager c300.

### Label retaining cell (LRC) protocol

The LRC protocol was carried out, as previously described. Briefly, wild-type mice received BrdU (7.5mg/ml) i.p injections twice daily for 5 days and sacrificed 4 wk after the last injection. Quantification of connexin protein colocalization with LRCs or ECs was performed on confocal z-stack images by a computational approach using FARSIGHT for nuclear segmentation and MATLAB programs as previously published in the lab^33^.

### Data analysis and statistics

All continuous variables were represented as mean ± SEM. The Mann-Whitney non-parametric test for unpaired samples was used to analyze continuous variables between groups and p value ≤ 0.05 was considered statistically significant. All analyses were performed using Prism 8.0 software (GraphPad).

---

**Extended Figure 1:**
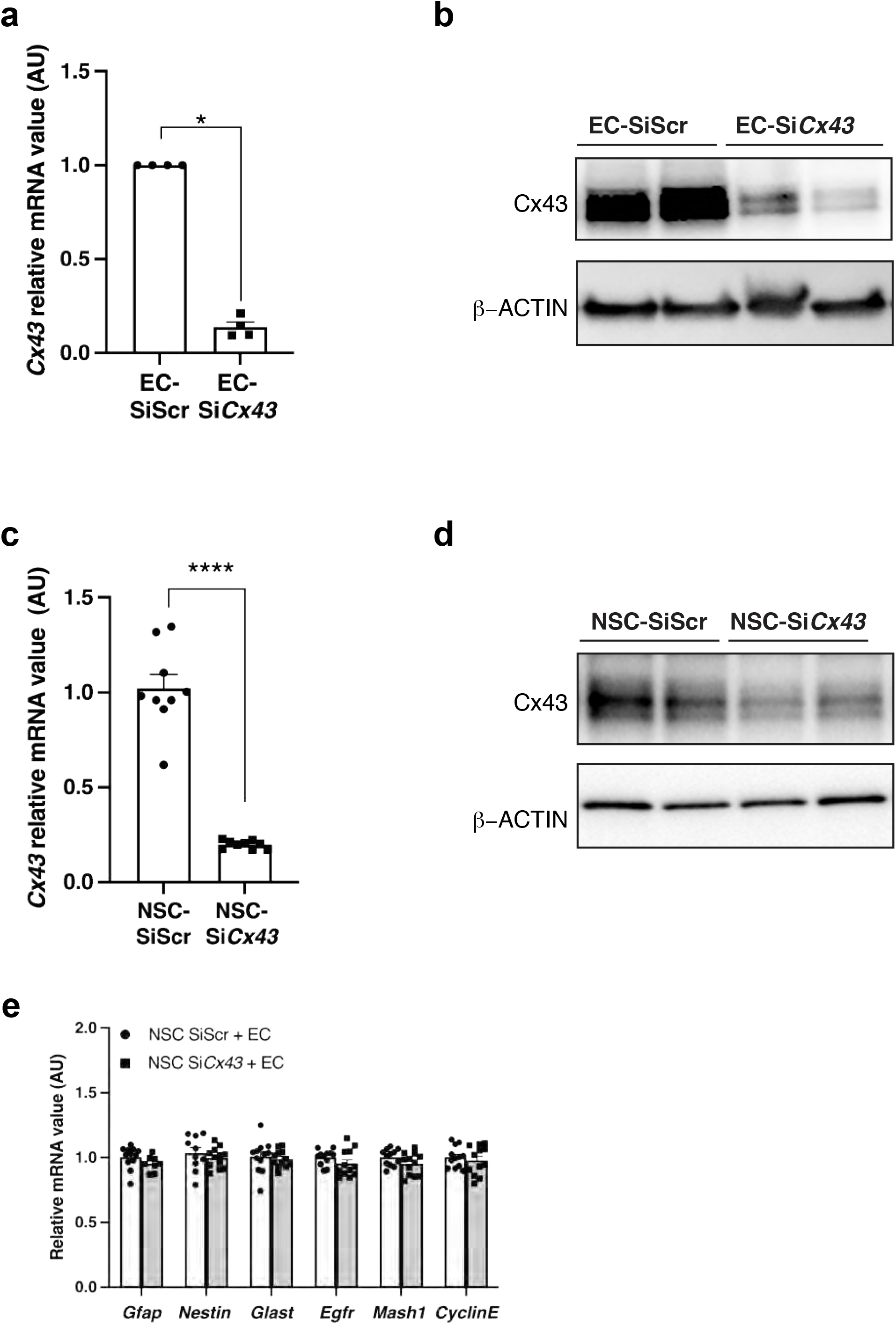
**(a)** qPCR analysis of *Cx43* mRNA level in EC treated with control siRNA (SiScr) or *Cx43* siRNA (si*Cx43*) (n=4 different experiments). (**b**) Representative western blot of Cx43 protein expression in EC treated with control siRNA (SiScr) or *Cx43* siRNA (si*Cx43*). **(c)** qPCR analysis of *Cx43* mRNA level in NSC treated with control siRNA (SiScr) or *Cx43* siRNA (si*Cx43*) (n=8 different experiments). (**d**) Representative western blot of Cx43 protein expression in NSC treated with control siRNA (SiScr) or *Cx43* siRNA (si*Cx43*). Data are mean ± SEM. \**p* ≤ 0.05, \*\*\*\**p* ≤ 0.0001. (**e**) qPCR analysis of quiescent and activated genes in NSC co-cultured with EC, where NSC are treated with control siRNA (siScr) or *Cx43* siRNA (Si*Cx43*) (n=3 different experiments).

**Extended Figure 2:**
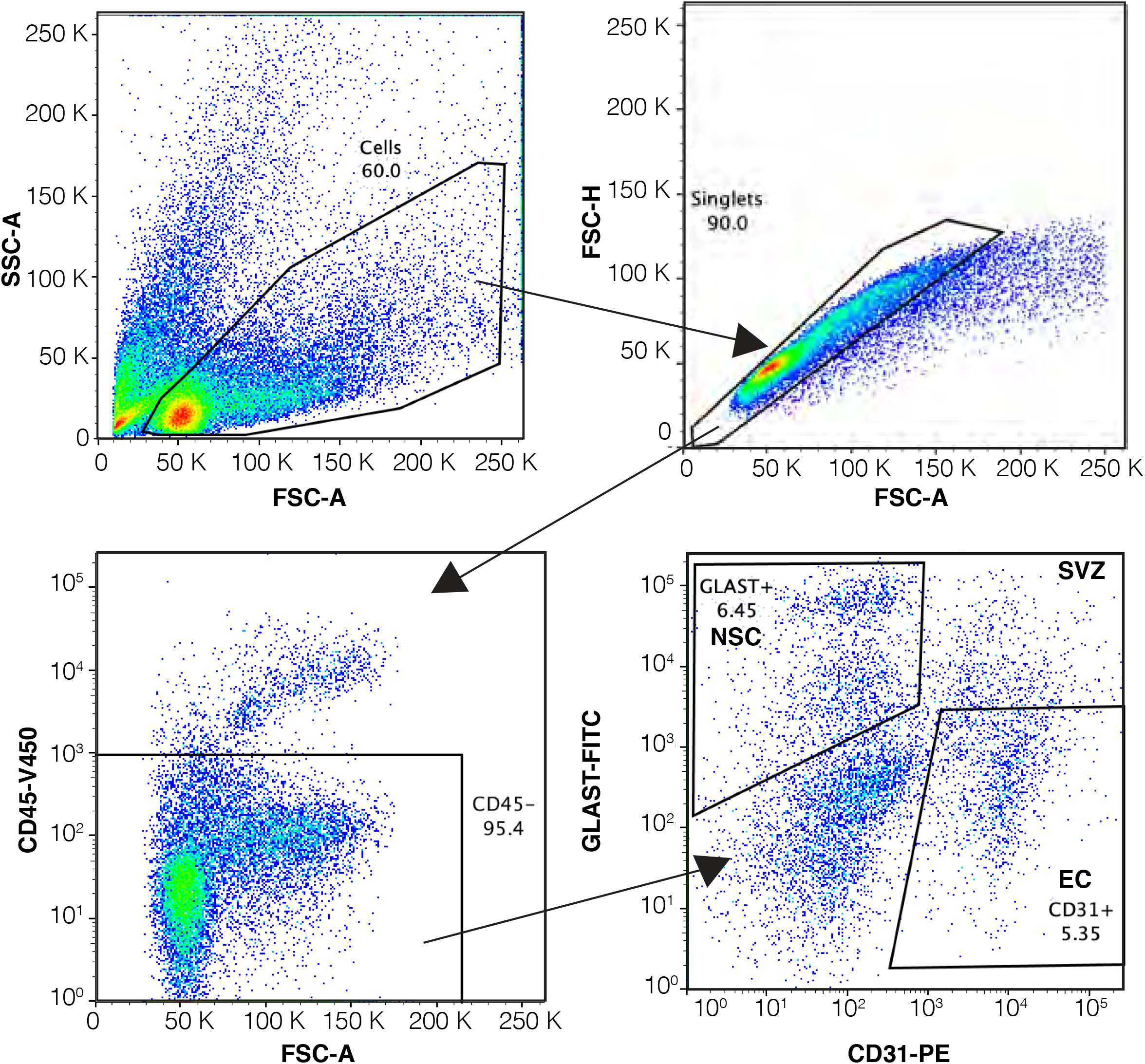
Representative FACS plot showing the gating strategy for ECs and NSCs isolation from adult mouse brain SVZ. After excluding debris (top left panel) and doublets (top right panel), CD45^-^ cells are selected (bottom left panel) and CD31^+^Glast^-^ NSC are collected (bottom right panel). Percentages refer to the population of cells in the previous parent gate. Plots show 100,000 events. SSA: Side-scatter area, FSC: forward-scatter area, FSH: forward-scatter height.

**Extended Figure 3:**
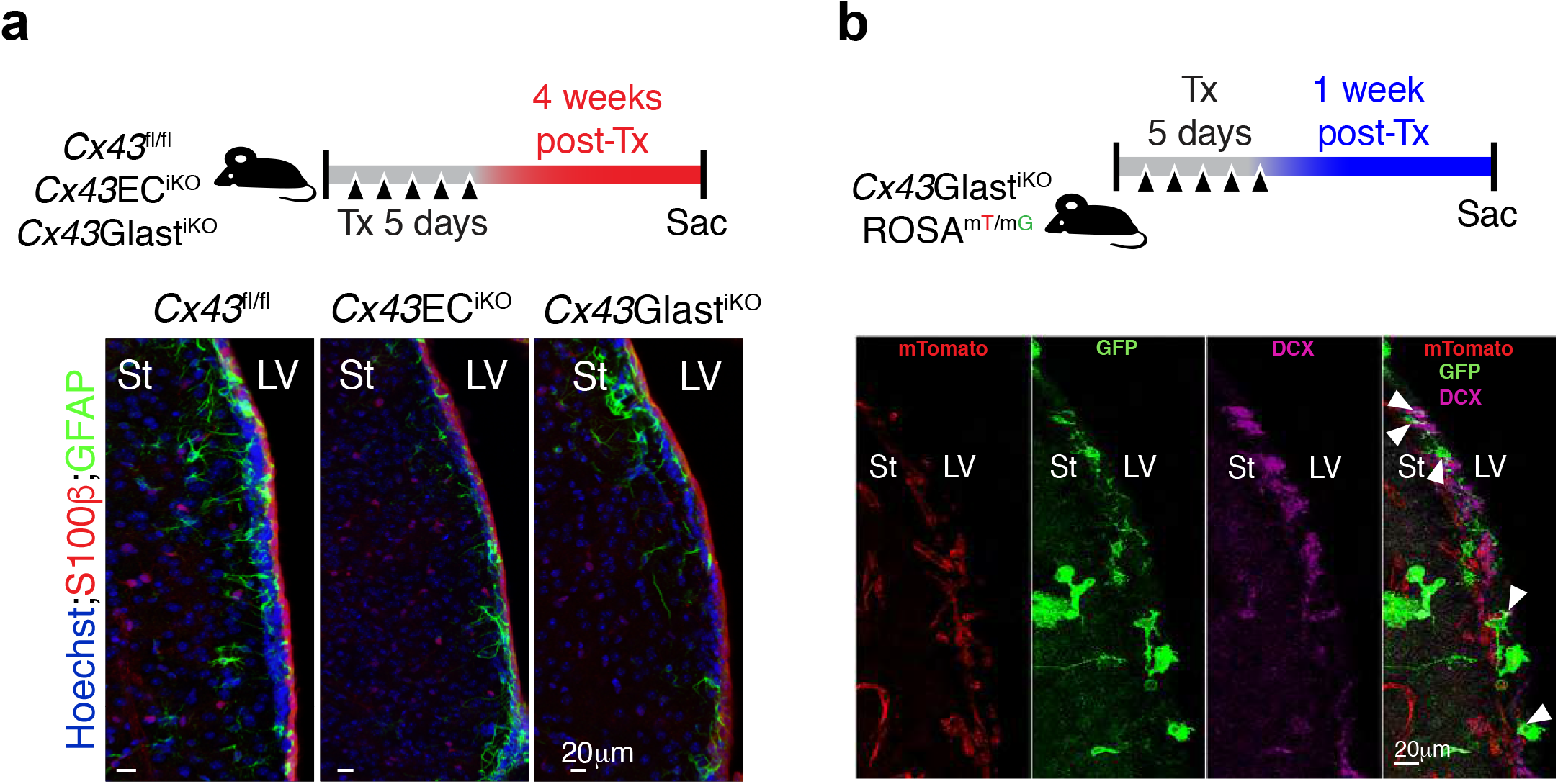
**(a)** Timeline used for label retaining cell (LRC) protocol in *Cx43*^fl/fl^, *Cx43*EC^iKO^ and *Cx43*Glast^iKO^ mice. (**b**) Representative confocal images of GFAP^+^SOX2^+^LRC in the SVZ of *Cx43*^fl/fl^, *Cx43*EC^iKO^ and *Cx43*Glast^iKO^ mice. (**c**) Quantification of images shown in (**b**) (n=10 different sections from 2 animals). (**d**) Timeline used for long-term *Cx43* recombination in *Cx43*^fl/fl^, *Cx43*EC^iKO^ and *Cx43*Glast^iKO^ mice. Bottom panel shows representative confocal images of S100β^+^GFAP^+^ cells in the SVZ of *Cx43*^fl/fl^, *Cx43*EC^iKO^ and *Cx43*Glast^iKO^ mice. (**e**) Timeline used for short-term *Cx43* recombination in *Cx43*Glast^iKO^;Rosa^mT/mG^ mice. Bottom panel shows a representative confocal image of a coronal SVZ section showing presence of DCX^+^GFP^+^ neuroblasts (white arrowheads) in the population of GFP^+^ recombinant cells. St: Striatum; LV: Lateral ventricle. Data are mean ± SEM. \*\**p* ≤ 0.01, \*\*\*\**p* ≤ 0.0001.

**Extended Figure 4:**
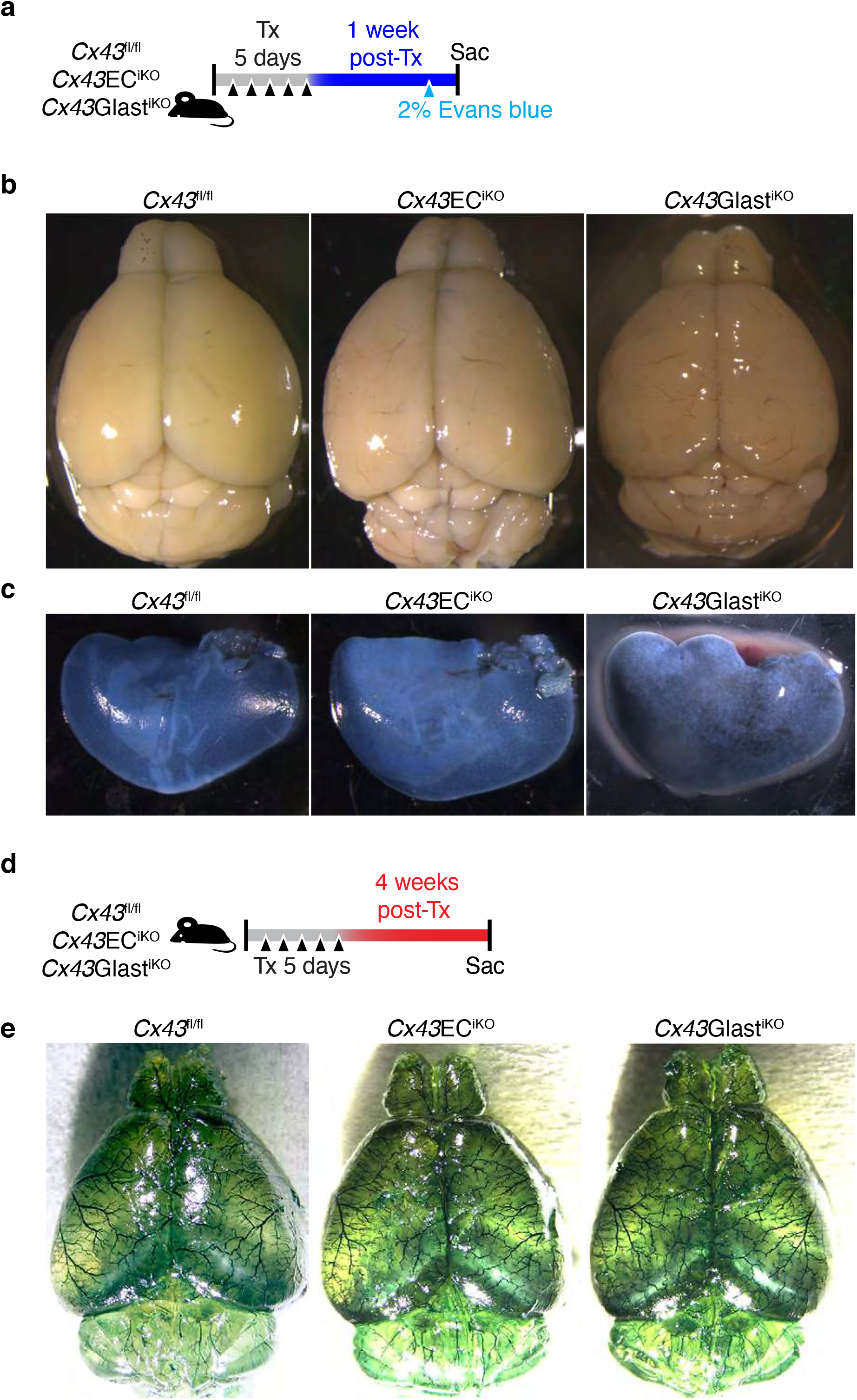
(**a**) Timeline used to evaluate Evans blue permeability in *Cx43*^fl/fl^, *Cx43*EC^iKO^ and *Cx43*Glast^iKO^ mice. (**b**) Representative brain images from *Cx4*3^fl/fl^, *Cx43*EC^iKO^ and *Cx43*Glast^iKO^ mice 24h post-2% Evans blue injection, showing absence of dye leakage. (**c**) Representative liver images from *Cx43*^fl/fl^, *Cx43*EC^iKO^ and *Cx43*Glast^iKO^ mice 24h post-2% Evans blue injection showing dye uptake. (**d**) Timeline used to visualize brain mircrovasculature network organization in *Cx43*^fl/fl^, *Cx43*EC^iKO^ and *Cx43*Glast^iKO^ mice. (**e**) Representative clarified brain images showing the microvasculature perfused with vascupaint green from *Cx4*3^fl/fl^, *Cx43*EC^iKO^ and *Cx43*Glast^iKO^.

**Extended Figure 5:**
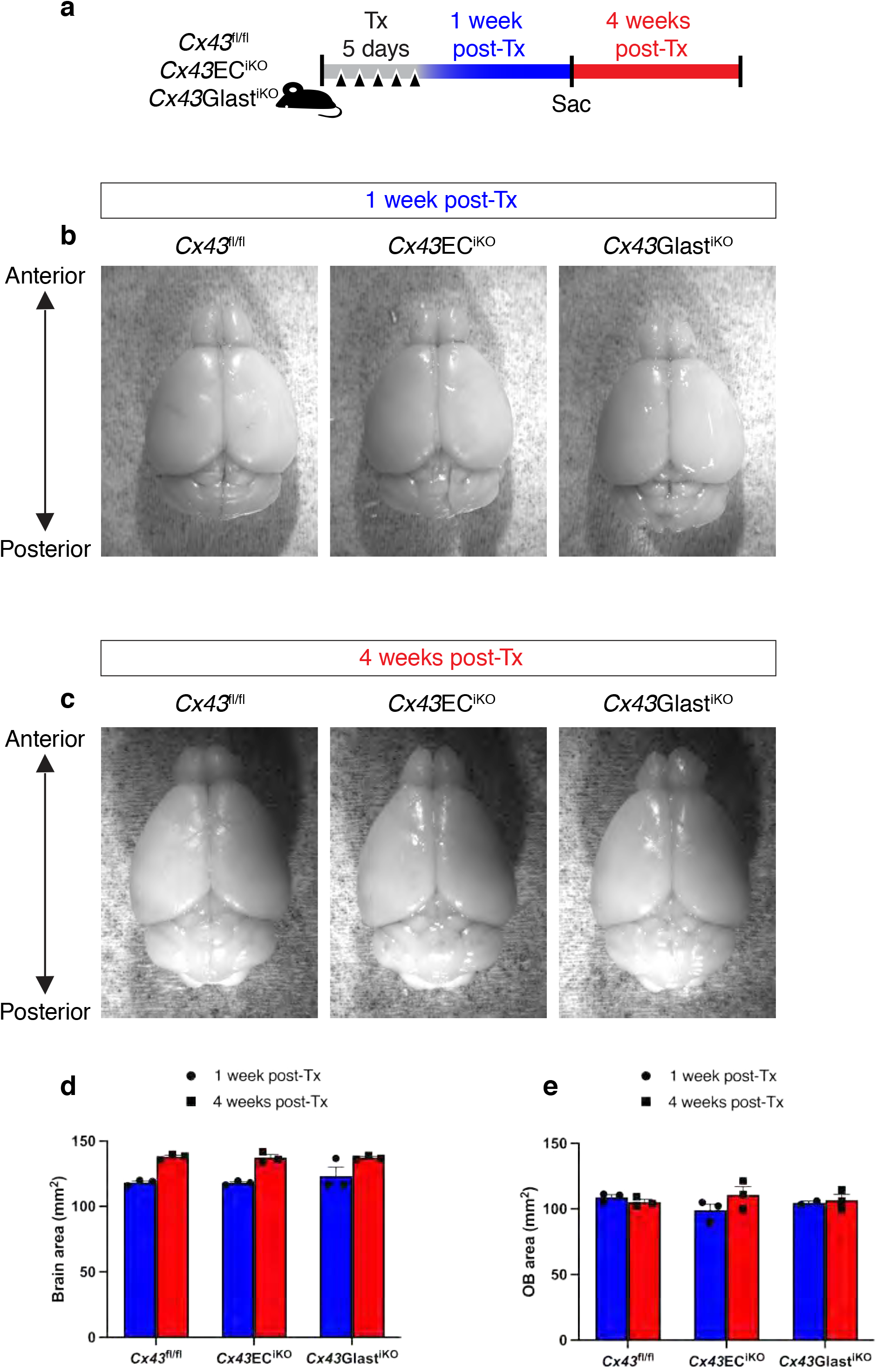
(**a**) Timeline used to evaluate brain and OB areas of *Cx43*^fl/fl^, *Cx43*EC^iKO^ and *Cx43*Glast^iKO^ mice. (**b**) and (**c**) Representative brain images from *Cx4*3^fl/fl^, *Cx43*EC^iKO^ and *Cx43*Glast^iKO^ mice 1-week and 4-weeks post-Tx injections respectively. (**d**) and (**e**) Bar graphs showing quantifications of brain and OB areas of *Cx43*^fl/fl^, *Cx43*EC^iKO^ and *Cx43*Glast^iKO^ mice respectively at 1-week and 4-weeks post-Tx injections.

**Extended Figure 6:**
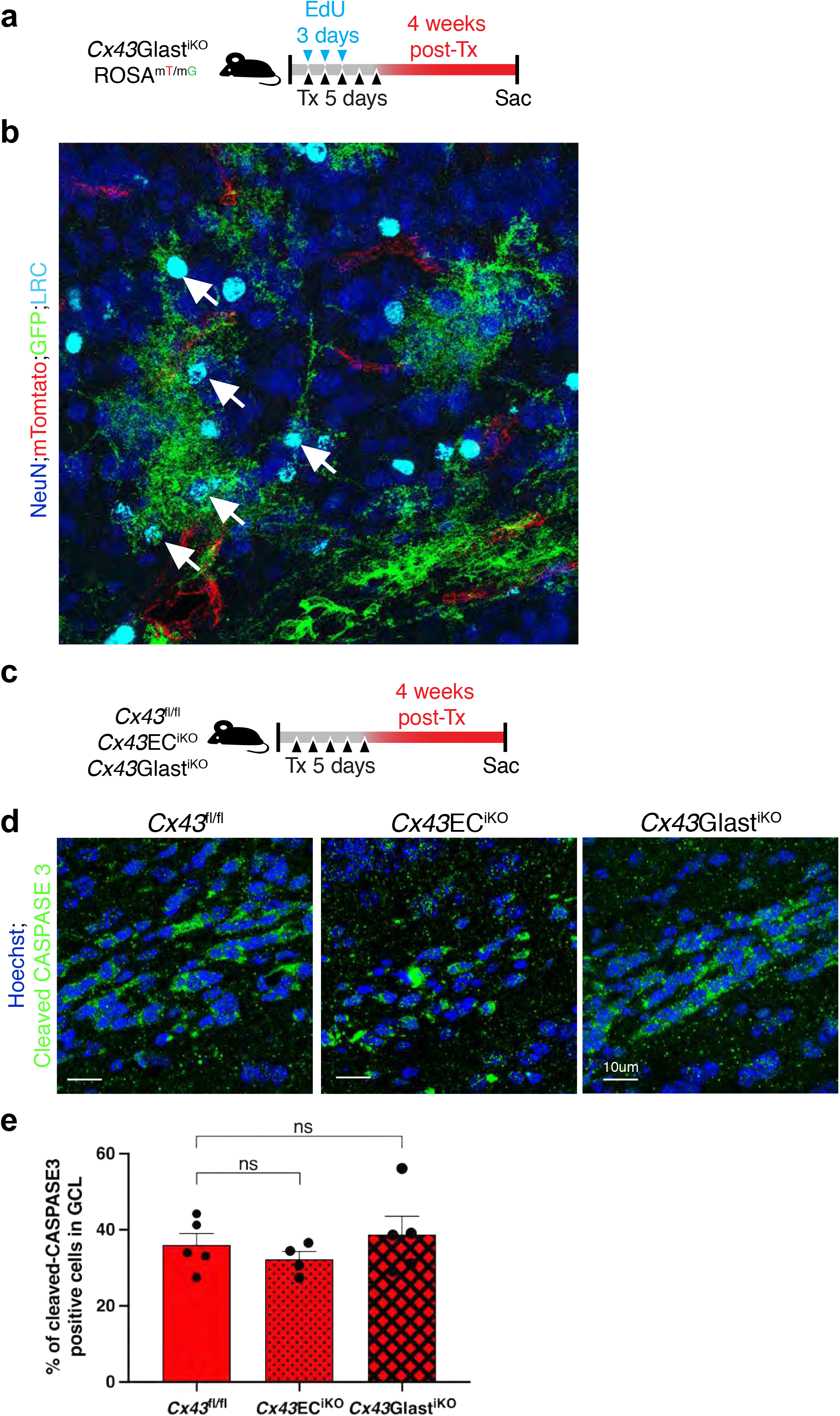
(**a**) Timeline used for *Cx43* recombination in *Cx43*Glast^iKO^;Rosa^mT/mG^ mice. (**b**) Representative confocal image of NeuN^+^LRC in the granule cell layer of the olfactory bulb of *Cx43*Glast^iKO^;Rosa^mT/mG^ mice. Note that NeuN^+^LRC are GFP^+^ (white arrows). (**c**) Timeline used for long-term *Cx43* recombination in *Cx43*^fl/fl^, *Cx43*EC^iKO^ and *Cx43*Glast^iKO^ mice. (**d**) Representative images of cleaved CASPASE-3^+^ cells in the granule cell layer of the olfactory bulb of *Cx43*^fl/fl^, *Cx43*EC^iKO^ and *Cx43*Glast^iKO^ mice. (**e**) Quantifications of images shown in (**d**) (n=4 different images analyzed from 1 animal per group). Data are mean ± SEM. ns > ≤ 0.05.

**Extended Figure 7:**
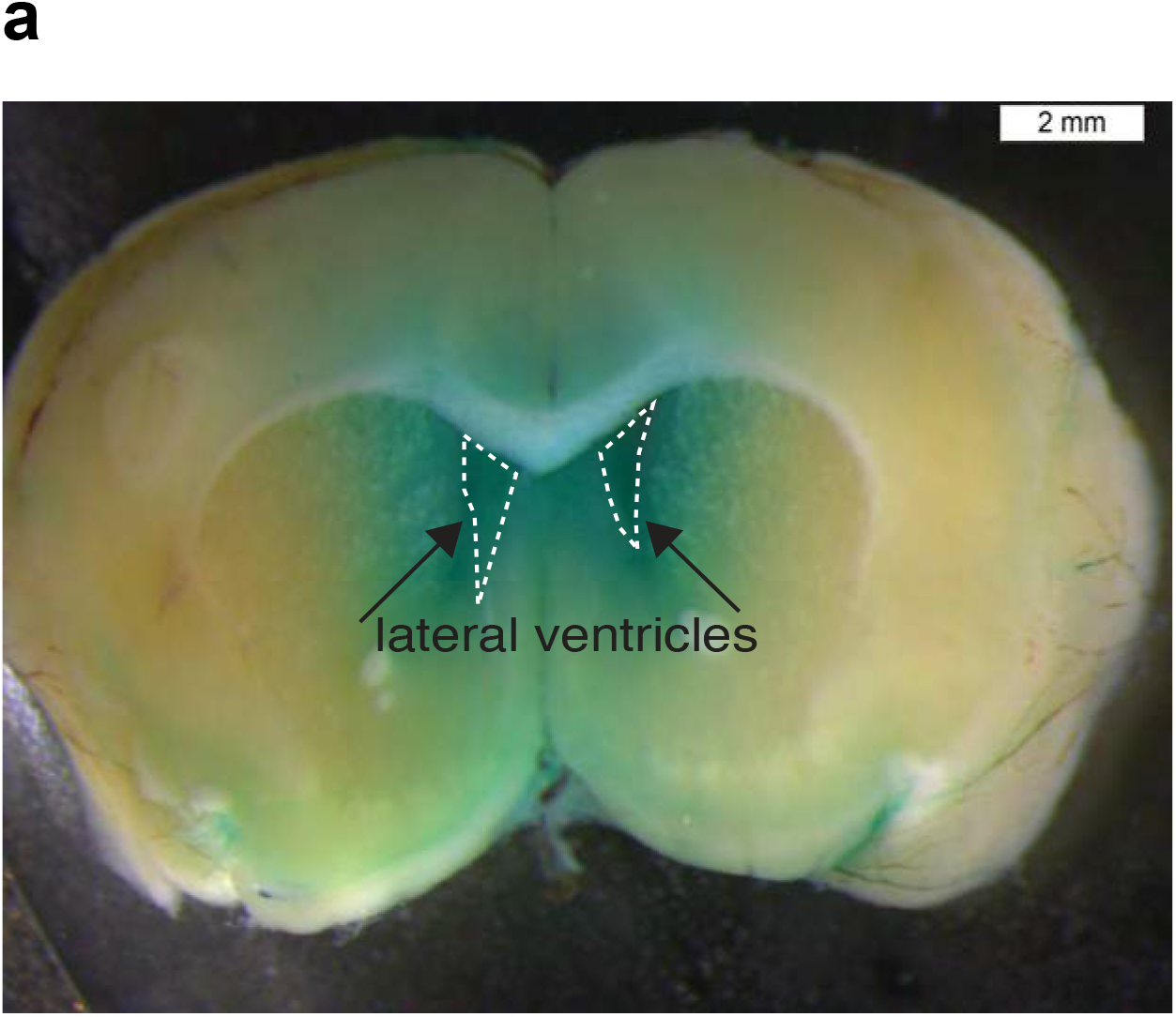
Image of brain sagittal section of a mouse injected with fast green dye in the ipsi-lateral ventricle at the optimized coordinates (x:1.4, y:0.5, z:2.5) used to infuse AraC. Note that both lateral ventricles (enclosed in dashed lines) are green colored after dye infusion, demonstrating that the coordinates used target the SVZ.

**Extended Figure 8:**
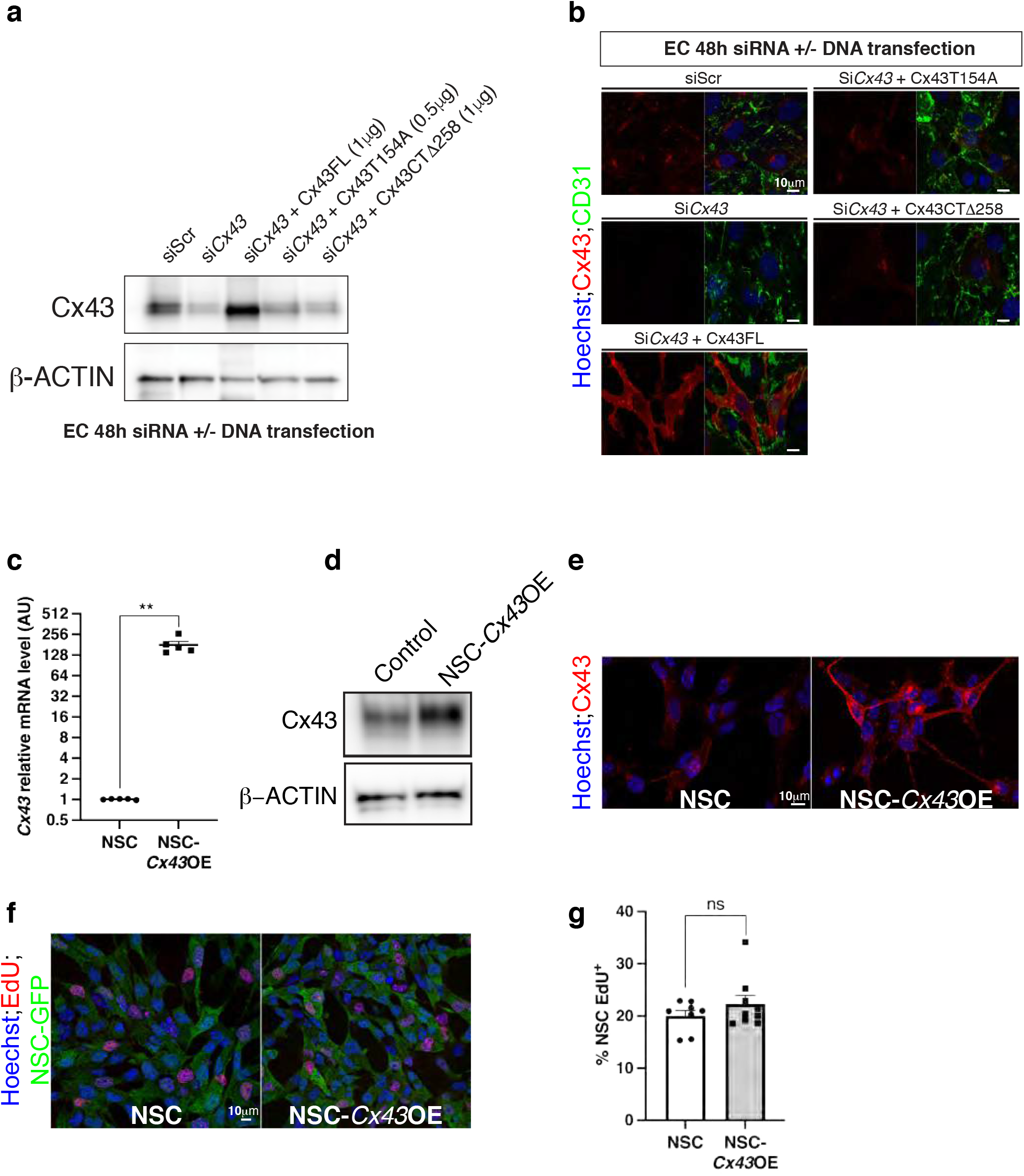
(**a**) Representative western blot of Cx43 protein expression in ECs treated with control siRNA (SiScr), *Cx43* siRNA (si*Cx43*), Si*Cx43* followed by Cx43 full length construct (FL), Si*Cx43* followed by Cx43T154A or Si*Cx43* followed by Cx43CTΔ258. (**b**) qPCR analysis of *Cx43* mRNA level in NSC and NSC overexpressing *Cx43* (NSC-*Cx43*OE) (n=5 different experiments). (**c**) Representative western blot of Cx43 protein expression in NSC and NSC-*Cx43OE.* (**d**) Representative confocal images of NSC and NSC-*Cx43*OE immuno-stained for Cx43. (**e**) Representative confocal images of NSC-GFP and NSC-GFP*-Cx43*OE 24h after EdU incorporation. (**f**) Quantification of images shown in (d) (n=4 different experiments). Data are mean ± SEM. ns > 0.05, \*\**p* ≤ 0.01.

## References

1 Chaker, Z., Codega, P. & Doetsch, F. A mosaic world: puzzles revealed by adult neural stem cell heterogeneity. Wiley Interdiscip Rev Dev Biol 5, 640–658, doi:10.1002/wdev.248 (2016).

2 Doetsch, F., Garcia-Verdugo, J. M. & Alvarez-Buylla, A. Cellular composition and three-dimensional organization of the subventricular germinal zone in the adult mammalian brain. J Neurosci 17, 5046–5061 (1997).

3 Parras, C. M. et al. Divergent functions of the proneural genes Mash1 and Ngn2 in the specification of neuronal subtype identity. Genes Dev 16, 324–338, doi:10.1101/gad.940902 (2002).

4 Pastrana, E., Cheng, L. C. & Doetsch, F. Simultaneous prospective purification of adult subventricular zone neural stem cells and their progeny. Proc Natl Acad Sci U S A 106, 6387–6392, doi:10.1073/pnas.0810407106 (2009).

5 Doetsch, F., Caille, I., Lim, D. A., Garcia-Verdugo, J. M. & Alvarez-Buylla, A. Subventricular zone astrocytes are neural stem cells in the adult mammalian brain. Cell 97, 703–716, doi:10.1016/s0092-8674(00)80783-7 (1999).

6 Doetsch, F., Garcia-Verdugo, J. M. & Alvarez-Buylla, A. Regeneration of a germinal layer in the adult mammalian brain. Proc Natl Acad Sci U S A 96, 11619–11624, doi:10.1073/pnas.96.20.11619 (1999).

7 Mirzadeh, Z., Merkle, F. T., Soriano-Navarro, M., Garcia-Verdugo, J. M. & Alvarez-Buylla, A. Neural stem cells confer unique pinwheel architecture to the ventricular surface in neurogenic regions of the adult brain. Cell Stem Cell 3, 265–278, doi:10.1016/j.stem.2008.07.004 (2008).

8 Gleeson, J. G., Lin, P. T., Flanagan, L. A. & Walsh, C. A. Doublecortin is a microtubule-associated protein and is expressed widely by migrating neurons. Neuron 23, 257–271, doi:10.1016/s0896-6273(00)80778-3 (1999).

9 Lois, C. & Alvarez-Buylla, A. Long-distance neuronal migration in the adult mammalian brain. Science 264, 1145–1148, doi:10.1126/science.8178174 (1994).

10 Menezes, J. R., Smith, C. M., Nelson, K. C. & Luskin, M. B. The division of neuronal progenitor cells during migration in the neonatal mammalian forebrain. Mol Cell Neurosci 6, 496–508, doi:10.1006/mcne.1995.0002 (1995).

11 Altman, J. Autoradiographic and histological studies of postnatal neurogenesis. IV. Cell proliferation and migration in the anterior forebrain, with special reference to persisting neurogenesis in the olfactory bulb. J Comp Neurol 137, 433–457, doi:10.1002/cne.901370404 (1969).

12 Luskin, M. B. Restricted proliferation and migration of postnatally generated neurons derived from the forebrain subventricular zone. Neuron 11, 173–189, doi:10.1016/0896-6273(93)90281-u (1993).

13 Corotto, F. S., Henegar, J. A. & Maruniak, J. A. Neurogenesis persists in the subependymal layer of the adult mouse brain. Neurosci Lett 149, 111–114, doi:10.1016/0304-3940(93)90748-a (1993).

14 James, R., Kim, Y., Hockberger, P. E. & Szele, F. G. Subventricular zone cell migration: lessons from quantitative two-photon microscopy. Front Neurosci 5, 30, doi:10.3389/fnins.2011.00030 (2011).

15 Obernier, K. et al. Adult Neurogenesis Is Sustained by Symmetric Self-Renewal and Differentiation. Cell Stem Cell 22, 221–234 e228, doi:10.1016/j.stem.2018.01.003 (2018).

16 Silva-Vargas, V., Delgado, A. C. & Doetsch, F. Symmetric Stem Cell Division at the Heart of Adult Neurogenesis. Neuron 98, 246–248, doi:10.1016/j.neuron.2018.04.005 (2018).

17 Shen, Q. et al. Adult SVZ stem cells lie in a vascular niche: a quantitative analysis of niche cell-cell interactions. Cell Stem Cell 3, 289–300, doi:10.1016/j.stem.2008.07.026 (2008).

18 Tavazoie, M. et al. A specialized vascular niche for adult neural stem cells. Cell Stem Cell 3, 279–288, doi:10.1016/j.stem.2008.07.025 (2008).

19 Calvo, C. F. et al. Vascular endothelial growth factor receptor 3 directly regulates murine neurogenesis. Genes Dev 25, 831–844, doi:10.1101/gad.615311 (2011).

20 Jin, K. et al. Vascular endothelial growth factor (VEGF) stimulates neurogenesis in vitro and in vivo. Proc Natl Acad Sci U S A 99, 11946–11950, doi:10.1073/pnas.182296499 (2002).

21 Gomez-Gaviro, M. V. et al. Betacellulin promotes cell proliferation in the neural stem cell niche and stimulates neurogenesis. Proc Natl Acad Sci U S A 109, 1317–1322, doi:10.1073/pnas.1016199109 (2012).

22 Ramirez-Castillejo, C. et al. Pigment epithelium-derived factor is a niche signal for neural stem cell renewal. Nat Neurosci 9, 331–339, doi:10.1038/nn1657 (2006).

23 Crouch, E. E., Liu, C., Silva-Vargas, V. & Doetsch, F. Regional and stage-specific effects of prospectively purified vascular cells on the adult V-SVZ neural stem cell lineage. J Neurosci 35, 4528–4539, doi:10.1523/JNEUROSCI.1188-14.2015 (2015).

24 Genet, N. & Hirschi, K. K. Understanding neural stem cell regulation in vivo and applying the insights to cell therapy for strokes. Regen Med 16, 861–870, doi:10.2217/rme-2021-0022 (2021).

25 Ottone, C. et al. Direct cell-cell contact with the vascular niche maintains quiescent neural stem cells. Nat Cell Biol 16, 1045–1056, doi:10.1038/ncb3045 (2014).

26 Bicker, F. et al. Neurovascular EGFL7 regulates adult neurogenesis in the subventricular zone and thereby affects olfactory perception. Nat Commun 8, 15922, doi:10.1038/ncomms15922 (2017).

27 Miragall, F., Albiez, P., Bartels, H., de Vries, U. & Dermietzel, R. Expression of the gap junction protein connexin43 in the subependymal layer and the rostral migratory stream of the mouse: evidence for an inverse correlation between intensity of connexin43 expression and cell proliferation activity. Cell Tissue Res 287, 243–253, doi:10.1007/s004410050749 (1997).

28 Nadarajah, B., Jones, A. M., Evans, W. H. & Parnavelas, J. G. Differential expression of connexins during neocortical development and neuronal circuit formation. J Neurosci 17, 3096–3111 (1997).

29 Bittman, K. S. & LoTurco, J. J. Differential regulation of connexin 26 and 43 in murine neocortical precursors. Cereb Cortex 9, 188–195, doi:10.1093/cercor/9.2.188 (1999).

30 Hirschi, K. K., Burt, J. M., Hirschi, K. D. & Dai, C. Gap junction communication mediates transforming growth factor-beta activation and endothelial-induced mural cell differentiation. Circ Res 93, 429–437, doi:10.1161/01.RES.0000091259.84556.D5 (2003).

31 Swayne, L. A. & Bennett, S. A. Connexins and pannexins in neuronal development and adult neurogenesis. BMC Cell Biol 17 Suppl 1, 10, doi:10.1186/s12860-016-0089-5 (2016).

32 Zhao, Y., Xin, Y., He, Z. & Hu, W. Function of Connexins in the Interaction between Glial and Vascular Cells in the Central Nervous System and Related Neurological Diseases. Neural Plast 2018, 6323901, doi:10.1155/2018/6323901 (2018).

33 Goldberg, J. S., Vadakkan, T. J., Hirschi, K. K. & Dickinson, M. E. A computational approach to detect gap junction plaques and associate them with cells in fluorescent images. J Histochem Cytochem 61, 283–293, doi:10.1369/0022155413477114 (2013).

34 Brazel, C. Y. et al. Sox2 expression defines a heterogeneous population of neurosphere-forming cells in the adult murine brain. Aging Cell 4, 197–207, doi:10.1111/j.1474-9726.2005.00158.x (2005).

35 Mercurio, S., Serra, L. & Nicolis, S. K. More than just Stem Cells: Functional Roles of the Transcription Factor Sox2 in Differentiated Glia and Neurons. Int J Mol Sci 20, doi:10.3390/ijms20184540 (2019).

36 Boulay, A. C. et al. Immune quiescence of the brain is set by astroglial connexin 43. J Neurosci 35, 4427–4439, doi:10.1523/JNEUROSCI.2575-14.2015 (2015).

37 Chew, S. S., Johnson, C. S., Green, C. R. & Danesh-Meyer, H. V. Role of connexin43 in central nervous system injury. Exp Neurol 225, 250–261, doi:10.1016/j.expneurol.2010.07.014 (2010).

38 Danesh-Meyer, H. V. & Green, C. R. Focus on molecules: connexin 43--mind the gap. Exp Eye Res 87, 494–495, doi:10.1016/j.exer.2008.01.021 (2008).

39 Kokovay, E. et al. VCAM1 is essential to maintain the structure of the SVZ niche and acts as an environmental sensor to regulate SVZ lineage progression. Cell Stem Cell 11, 220–230, doi:10.1016/j.stem.2012.06.016 (2012).

40 DeVos, S. L. & Miller, T. M. Direct intraventricular delivery of drugs to the rodent central nervous system. J Vis Exp, e50326, doi:10.3791/50326 (2013).

41 Beahm, D. L. et al. Mutation of a conserved threonine in the third transmembrane helix of alpha- and beta-connexins creates a dominant-negative closed gap junction channel. J Biol Chem 281, 7994–8009, doi:10.1074/jbc.M506533200 (2006).

42 Maass, K., Shibayama, J., Chase, S. E., Willecke, K. & Delmar, M. C-terminal truncation of connexin43 changes number, size, and localization of cardiac gap junction plaques. Circ Res 101, 1283–1291, doi:10.1161/CIRCRESAHA.107.162818 (2007).

43 Genet, N., Bhatt, N., Bourdieu, A. & Hirschi, K. K. Multifaceted Roles of Connexin 43 in Stem Cell Niches. Curr Stem Cell Rep 4, 1–12, doi:10.1007/s40778-018-0110-3 (2018).

44 Moorer, M. C. et al. Defective signaling, osteoblastogenesis and bone remodeling in a mouse model of connexin 43 C-terminal truncation. J Cell Sci 130, 531–540, doi:10.1242/jcs.197285 (2017).

45 Andreu-Agullo, C., Morante-Redolat, J. M., Delgado, A. C. & Farinas, I. Vascular niche factor PEDF modulates Notch-dependent stemness in the adult subependymal zone. Nat Neurosci 12, 1514–1523, doi:10.1038/nn.2437 (2009).

46 Kirschenbaum, B. & Goldman, S. A. Brain-derived neurotrophic factor promotes the survival of neurons arising from the adult rat forebrain subependymal zone. Proc Natl Acad Sci U S A 92, 210–214, doi:10.1073/pnas.92.1.210 (1995).

47 Kokovay, E. et al. Adult SVZ lineage cells home to and leave the vascular niche via differential responses to SDF1/CXCR4 signaling. Cell Stem Cell 7, 163–173, doi:10.1016/j.stem.2010.05.019 (2010).

48 Leventhal, C., Rafii, S., Rafii, D., Shahar, A. & Goldman, S. A. Endothelial trophic support of neuronal production and recruitment from the adult mammalian subependyma. Mol Cell Neurosci 13, 450–464, doi:10.1006/mcne.1999.0762 (1999).

49 Snapyan, M. et al. Vasculature guides migrating neuronal precursors in the adult mammalian forebrain via brain-derived neurotrophic factor signaling. J Neurosci 29, 4172–4188, doi:10.1523/JNEUROSCI.4956-08.2009 (2009).

50 Cheng, A. et al. Gap junctional communication is required to maintain mouse cortical neural progenitor cells in a proliferative state. Dev Biol 272, 203–216, doi:10.1016/j.ydbio.2004.04.031 (2004).

51 Duval, N. et al. Cell coupling and Cx43 expression in embryonic mouse neural progenitor cells. J Cell Sci 115, 3241–3251 (2002).

52 Ravella, A., Ringstedt, T., Brion, J. P., Pandolfo, M. & Herlenius, E. Adult neural precursor cells form connexin-dependent networks that improve their survival. Neuroreport 26, 928–936, doi:10.1097/WNR.0000000000000451 (2015).

53 Todorova, M. G., Soria, B. & Quesada, I. Gap junctional intercellular communication is required to maintain embryonic stem cells in a non-differentiated and proliferative state. J Cell Physiol 214, 354–362, doi:10.1002/jcp.21203 (2008).

54 Gairhe, S., Bauer, N. N., Gebb, S. A. & McMurtry, I. F. Myoendothelial gap junctional signaling induces differentiation of pulmonary arterial smooth muscle cells. Am J Physiol Lung Cell Mol Physiol 301, L527–535, doi:10.1152/ajplung.00091.2011 (2011).

55 Isakson, B. E. & Duling, B. R. Heterocellular contact at the myoendothelial junction influences gap junction organization. Circ Res 97, 44–51, doi:10.1161/01.RES.0000173461.36221.2e (2005).

56 Matta, R. et al. Minimally Invasive Delivery of Microbeads with Encapsulated, Viable and Quiescent Neural Stem Cells to the Adult Subventricular Zone. Sci Rep 9, 17798, doi:10.1038/s41598-019-54167-1 (2019).

57 Warn-Cramer, B. J. et al. Characterization of the mitogen-activated protein kinase phosphorylation sites on the connexin-43 gap junction protein. J Biol Chem 271, 3779–3786, doi:10.1074/jbc.271.7.3779 (1996).

58 Solan, J. L., Marquez-Rosado, L. & Lampe, P. D. Cx43 phosphorylation-mediated effects on ERK and Akt protect against ischemia reperfusion injury and alter the stability of the stress-inducible protein NDRG1. J Biol Chem 294, 11762–11771, doi:10.1074/jbc.RA119.009162 (2019).

59 Sorensen, I., Adams, R. H. & Gossler, A. DLL1-mediated Notch activation regulates endothelial identity in mouse fetal arteries. Blood 113, 5680–5688, doi:10.1182/blood-2008-08-174508 (2009).

60 Wang, Y. et al. Ephrin-B2 controls VEGF-induced angiogenesis and lymphangiogenesis. Nature 465, 483–486, doi:10.1038/nature09002 (2010).

61 Mori, T. et al. Inducible gene deletion in astroglia and radial glia--a valuable tool for functional and lineage analysis. Glia 54, 21–34, doi:10.1002/glia.20350 (2006).

62 Cotsarelis, G., Cheng, S. Z., Dong, G., Sun, T. T. & Lavker, R. M. Existence of slow-cycling limbal epithelial basal cells that can be preferentially stimulated to proliferate: implications on epithelial stem cells. Cell 57, 201–209, doi:10.1016/0092-8674(89)90958-6 (1989).

63 Potten, C. S. & Morris, R. J. Epithelial stem cells in vivo. J Cell Sci Suppl 10, 45–62, doi:10.1242/jcs.1988.supplement_10.4 (1988).

64 Kazanis, I. et al. Quiescence and activation of stem and precursor cell populations in the subependymal zone of the mammalian brain are associated with distinct cellular and extracellular matrix signals. J Neurosci 30, 9771–9781, doi:10.1523/JNEUROSCI.0700-10.2010 (2010).

65 Crouch, E. E. & Doetsch, F. FACS isolation of endothelial cells and pericytes from mouse brain microregions. Nat Protoc 13, 738–751, doi:10.1038/nprot.2017.158 (2018).

66 Livak, K. J. & Schmittgen, T. D. Analysis of relative gene expression data using real-time quantitative PCR and the 2(-Delta Delta C(T)) Method. Methods 25, 402–408, doi:10.1006/meth.2001.1262 (2001).

67 Pollard, S. M. In vitro expansion of fetal neural progenitors as adherent cell lines. Methods Mol Biol 1059, 13–24, doi:10.1007/978-1-62703-574-3_2 (2013).

68 Conti, L. et al. Niche-independent symmetrical self-renewal of a mammalian tissue stem cell. PLoS Biol 3, e283, doi:10.1371/journal.pbio.0030283 (2005).

69 Pollard, S. M. & Conti, L. Investigating radial glia in vitro. Prog Neurobiol 83, 53–67, doi:10.1016/j.pneurobio.2007.02.008 (2007).

70 Afgan, E. et al. The Galaxy platform for accessible, reproducible and collaborative biomedical analyses: 2018 update. Nucleic Acids Res 46, W537–W544, doi:10.1093/nar/gky379 (2018).

71 Blankenberg, D. et al. Manipulation of FASTQ data with Galaxy. Bioinformatics 26, 1783–1785, doi:10.1093/bioinformatics/btq281 (2010).

72 Blankenberg, D. et al. Galaxy: a web-based genome analysis tool for experimentalists. Curr Protoc Mol Biol Chapter 19, Unit 19 10 11–21, doi:10.1002/0471142727.mb1910s89 (2010).

73 Bray, N. L., Pimentel, H., Melsted, P. & Pachter, L. Near-optimal probabilistic RNA-seq quantification. Nat Biotechnol 34, 525–527, doi:10.1038/nbt.3519 (2016).

74 Pimentel, H., Bray, N. L., Puente, S., Melsted, P. & Pachter, L. Differential analysis of RNA-seq incorporating quantification uncertainty. Nat Methods 14, 687–690, doi:10.1038/nmeth.4324 (2017).

